# Mitochondrial transplantation redirects rather than suppresses the inflammatory response in primary human monocytes

**DOI:** 10.64898/2026.11.06.762873

**Authors:** Saeed Nazemidashtarjandi, Sairam Pantham, Rachel Dori, Dong Bin Back, Gen Hamanaka, Kazuhide Hawakaya, Martin L. Yarmush

## Abstract

Circulating monocytes are recruited into inflamed tissue, where local cues determine whether the macrophages they become drive inflammation or resolve it. Because that decision occurs after cells leave the blood, the circulating stage offers a window to alter monocyte programming, and monocyte inflammatory function is closely coupled to mitochondrial metabolism. Here we transplanted freshly isolated, respiration-competent mitochondria from HEK293T cells into primary human CD14⁺ monocytes and characterized the resulting mitochondria-transplanted monocytes (MTM) across surface phenotype, respiration, and inflammatory function. Isolated mitochondria were intact and respiration-competent, with a respiratory control ratio of 8.7 ± 2.2 and a coupling efficiency of 78.9 ± 8.5%. Monocytes internalized them in proportion to the dose supplied; within 3 h, 88% of viable cells were positive for labeled mitochondria, with no detected loss of viability; isolated mitochondria did not activate NF-κB or IRF reporters in THP1-Dual cells. Transplantation reduced the frequency of the pro-inflammatory intermediate subset, raised CD14 2.11-fold and HLA-DR 1.31-fold, and lowered CD163. MTM showed reduced ATP-linked respiration with maximal capacity preserved, and a trend toward a higher maximal-to-basal respiration ratio. Following stimulation with LPS, MTM released markedly less TNF-α (0.06-fold) and CXCL10/IP-10 (0.25-fold) and, in most donors, more IL-10 (3.33-fold), while the remaining analytes were largely unchanged. Mitochondrial transplantation therefore alters primary human monocytes toward a phenotype with selectively reduced inflammatory mediator release and increased IL-10 in most donors and identifies circulating monocytes as a tractable target for mitochondria-based modulation of inflammation.

**Translational Impact Statement:** Monocytes are an overlooked therapeutic target. They circulate briefly, then extravasate into inflamed tissue and become the macrophages that sustain or resolve it, so conditioning them before they arrive acts upstream of the damage rather than after it. We show that transferring functional mitochondria into primary human monocytes selectively dampens their endotoxin response, lowering TNF-α and CXCL10 while raising IL-10 in most donors. Mitochondrial transfer therefore offers a way to redirect these cells before they reach the tissue they will act in.

## Introduction

Human blood monocytes comprise classical (CD14++CD16−), intermediate (CD14++CD16+) and non-classical (CD14+CD16++) populations.^1,2^ In vivo labeling in healthy volunteers shows that classical monocytes dominate the circulating pool and differentiate sequentially into intermediate and non-classical cells rather than committing to a tissue fate.^3^ After recruitment into inflamed tissue, monocytes encounter local growth factors, cytokines and microbial products that determine whether the macrophages and dendritic cells they become drive inflammation or resolve it.^4^ Because human and mouse subsets correspond only approximately,^5^ this behavior is best tested in human cells.

Monocyte differentiation remains responsive to local signals after recruitment, and circulating monocytes are accessible without delivery into inflamed tissue, which is why the monocyte-derived compartment has been proposed as amenable to therapeutic manipulation.^4^ The intestine is the clearest illustration: intestinal macrophages turn over rapidly and are continuously replenished from blood monocytes throughout adult life,^4,6^ and where their maturation is disturbed, as in inflammatory bowel disease (IBD), pro-inflammatory monocyte-like cells accumulate in the mucosa in place of tolerogenic resident macrophages.^7–9^ These changes are not confined to the mucosa: the intermediate CD14++CD16+ subset is expanded in the blood of patients with active Crohn’s disease,^10^ and inflammatory programming present in circulating monocytes is retained by their intestinal descendants.^11^ Whether that balance can be deliberately altered before tissue recruitment is largely untested.

Monocyte inflammatory function is coupled to metabolic state. Microbial stimulation of human monocytes induces distinct metabolic programs depending on the receptor engaged, with LPS driving a pronounced increase in glycolytic flux.^12^ In mononuclear phagocytes, mitochondria are repurposed from ATP synthesis toward signaling intermediates that amplify cytokine output,^13,14^ whereas oxidative capacity is required for the resolving arm of the response, and mitochondrial dysfunction prevents inflammatory macrophages from repolarizing.^15^

The link runs in both directions through IL-10, which blocks LPS-induced metabolic rewiring by inducing DDIT4 and inhibiting mTOR, suppressing glycolysis and promoting oxidative phosphorylation; where IL-10 signaling is lost, damaged mitochondria accumulate in macrophages and drive NLRP3 inflammasome activation and IL-1β production, as observed in experimental colitis and in patients with IBD.^16^ Mitochondrial capacity is therefore part of the machinery that sets the inflammatory program, and it is modifiable in human disease: circulating monocytes from patients in septic shock show impaired bioenergetic function that recovers with clinical improvement.^17,18^

Mitochondria can be transferred between cells and internalized as intact, functional organelles. Intercellular transfer occurs between mammalian cells in culture^19^ and in vivo from mesenchymal stromal cells to injured alveolar epithelium,^20^ from astrocytes to neurons,^21^ and from adipocytes to tissue macrophages, where it regulates macrophage function and is impaired in obesity.^22^ This biology has been engineered into a therapeutic intervention: autologous mitochondria delivered to ischemic myocardium are internalized through actin-dependent macropinocytosis^23^ and improve post-ischemic contractile function,^24^ an approach since applied in patients.^25^

Mitochondrial transplantation has therefore been investigated extensively in parenchymal tissues, but its effects on primary human monocytes remain incompletely characterized. Exogenous mitochondrial delivery attenuates LPS-induced inflammatory signalling in mouse alveolar macrophages,^26^ indicating that myeloid cells are responsive, but whether primary human monocytes internalize transplanted mitochondria, and whether doing so alters their surface phenotype, respiratory capacity and inflammatory output, is unresolved.

Here we transplanted freshly isolated, respiration-competent mitochondria into primary human CD14+ monocytes and characterized the resulting mitochondria-transplanted monocytes (MTM) across surface phenotype, respiration and inflammatory function, asking whether conditioning monocytes in the circulating state alters the response they mount when subsequently challenged with LPS.

## Results

### Isolated mitochondria are intact and respiration-competent

To examine how mitochondrial transplantation alters human monocytes, we isolated CD14⁺ monocytes from human peripheral blood by density-gradient centrifugation followed by positive magnetic selection, and prepared mitochondria from HEK293T cells by differential centrifugation (**Figure 1A**).

**Figure 1.**
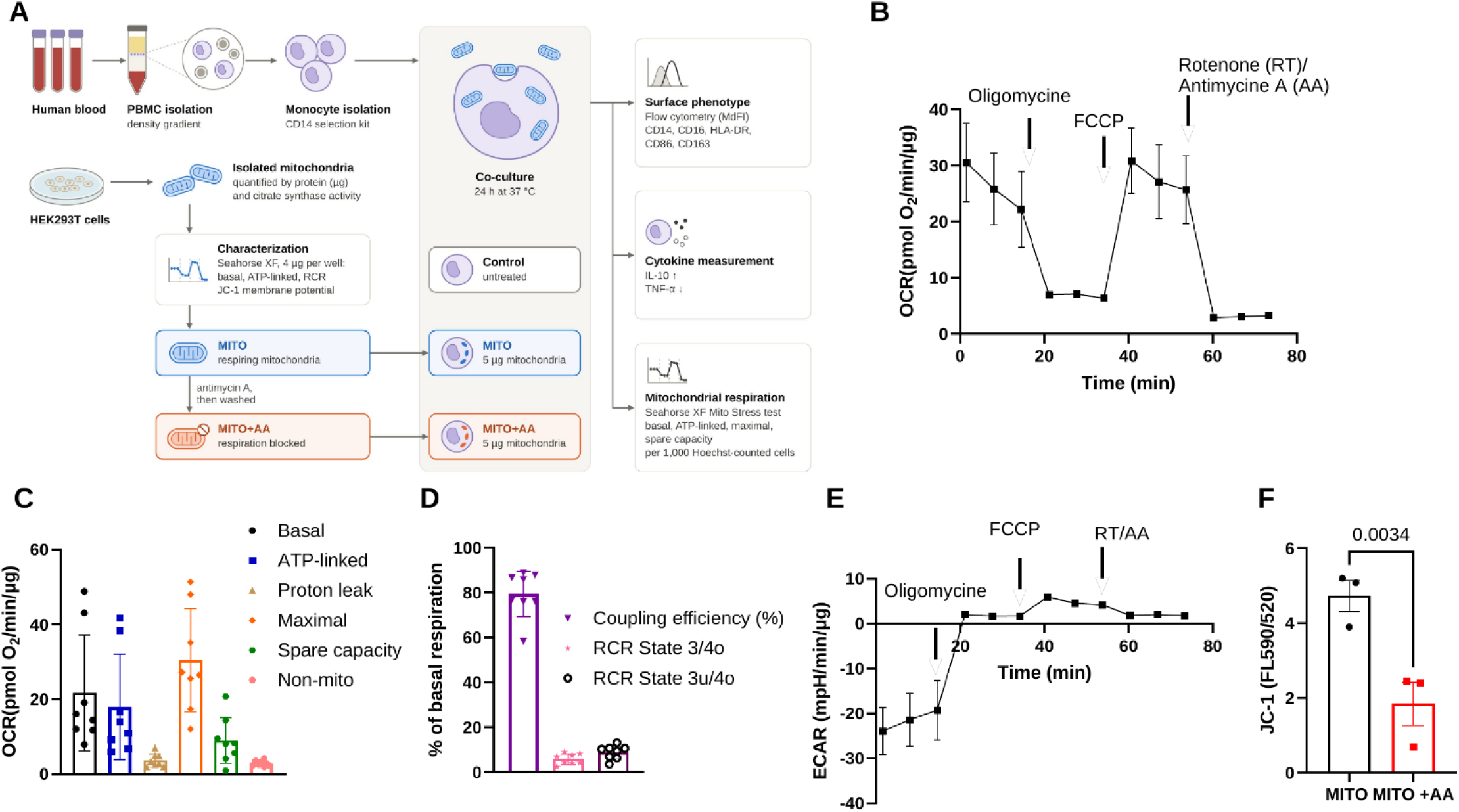
Study design and characterization of isolated mitochondria. (A) Experimental workflow. Peripheral blood mononuclear cells (PBMCs) were isolated from human blood by density-gradient centrifugation, and monocytes were purified by CD14-positive selection. Mitochondria were isolated from HEK293T cells and quantified by protein content and citrate synthase activity. An aliquot was treated with antimycin A (AA) and washed to give respiration-deficient mitochondria (MITO+AA). Monocytes were cultured for 24 h at 37 °C alone (Control) or with 5 µg of respiration-competent (MITO) or antimycin A-treated (MITO+AA) mitochondrial protein, then analyzed for monocyte subset and activation markers (CD14, CD16, HLA-DR, CD86, CD163) by flow cytometry, for phagocytosis of pHrodo-labeled particles, for mitochondrial respiration (Seahorse XF Cell Mito Stress Test), and for IL-10 and TNF-α release after LPS stimulation. (B) Oxygen consumption rate (OCR) of isolated mitochondria at 4 µg of mitochondrial protein per well, in mitochondrial assay solution containing pyruvate, malate, and ADP, with sequential injection of oligomycin, FCCP, and rotenone/antimycin A. (C) Derived respiratory parameters, basal (ADP-stimulated, State 3), ATP-linked, proton leak, maximal, spare capacity, and residual oxygen consumption after rotenone/antimycin A, each expressed per µg of mitochondrial protein. (D) Coupling efficiency and respiratory control ratio, calculated both from the uncoupled rate (State 3u/State 4o) and from the ADP-stimulated rate (State 3/State 4o). (E) Extracellular acidification rate (ECAR) over the same injection sequence; the negative basal rate indicates net alkalinization of the medium in the absence of glycolysis. (F) Mitochondrial membrane potential measured with JC-1 and expressed as the ratio of aggregate to monomer fluorescence (in MITO and MITO+AA. Data in (B–E) are mean ± SEM of n = 5 independent mitochondrial preparations; the effect of each injection in (B) was tested by a paired t-test with Holm–Šidák correction. (F) Mean ± SEM of three independent preparations, compared by paired t-test.

Respiration was measured with the Seahorse XF Cell Mito Stress Test in mitochondrial assay solution containing pyruvate, malate, and ADP, so that basal respiration corresponds to the ADP-stimulated rate (State 3) rather than a resting rate (**Figure 1B**). Across five independent preparations at 4 µg of mitochondrial protein per well, oligomycin reduced OCR to 0.32-fold of basal (P = 0.003), FCCP restored it to 4.71-fold of the oligomycin-inhibited rate (P < 0.001), and rotenone with antimycin A reduced it to 0.095-fold of the FCCP-stimulated rate (P < 0.001) (repeated-measures one-way ANOVA on log OCR, P < 0.001 with Geisser-Greenhouse correction). Rotenone and antimycin A abolished 83.4 ± 8.3% of basal oxygen consumption, confirming that most of the signal was respiratory-chain dependent.

**Figure 1C** shows the derived parameters. Basal respiration was 19.3 ± 15.3, ATP-linked respiration 15.9 ± 13.9, proton leak 3.41 ± 1.64, maximal respiration 28.2 ± 13.5, and residual oxygen consumption after rotenone/antimycin A 2.81 ± 0.68 pmol O₂/min per µg of mitochondrial protein (mean ± SD, n = 5). Coupling efficiency was 78.9 ± 8.5%, and the respiratory control ratio, uncoupled maximal respiration (State 3u) divided by oligomycin-insensitive leak respiration (State 4o), was 8.7 ± 2.2 (**Figure 1D**), consistent with intact inner membranes. Because basal respiration was already ADP-stimulated, the measured spare capacity (8.8 ± 5.0 pmol O₂/min/µg) reflects the margin between State 3 and uncoupled respiration rather than a resting reserve.

Extracellular acidification behaved as expected for isolated organelles rather than intact cells (**Figure 1E**). Basal ECAR was negative (−19.3 ± 14.9 mpH/min), indicating net alkalinization of the medium. Oligomycin abolished the alkalinization (ECAR +1.87 ± 0.56 mpH/min; shift of 21.2 ± 14.6 mpH/min), and the shift size tracked ATP-linked respiration across preparations (Pearson r = 0.997, P < 0.001, n = 5), an association that does not establish the underlying proton flux. Finally, mitochondrial membrane potential measured with JC-1 confirmed that the preparations were energized, and inhibition of the electron transport chain with antimycin A (AA) reduced the potential (MITO vs MITO+AA, n = 3; P = 0.0034, paired t-test; **Figure 1F**).

### Human monocytes take up isolated mitochondria

We next asked whether circulating human monocytes internalize isolated mitochondria. Citrate synthase activity of the preparations increased linearly with mitochondrial protein across the range used (P = 0.007; **Supplementary Figure 1**), supporting protein mass as a proxy for mitochondrial content in these preparations. Doses are therefore given as micrograms of mitochondrial protein throughout.

Monocytes were co-cultured for 24 h at 37 °C with isolated mitochondria at 5, 10, 15 or 25 µg of mitochondrial protein, and activity was then measured in the washed monocytes. Citrate synthase activity increased with the transplanted dose (**Figure 2A**; n = 3), consistent with a dose-dependent increase in monocyte-associated mitochondrial enzyme activity.

**Figure 2.**
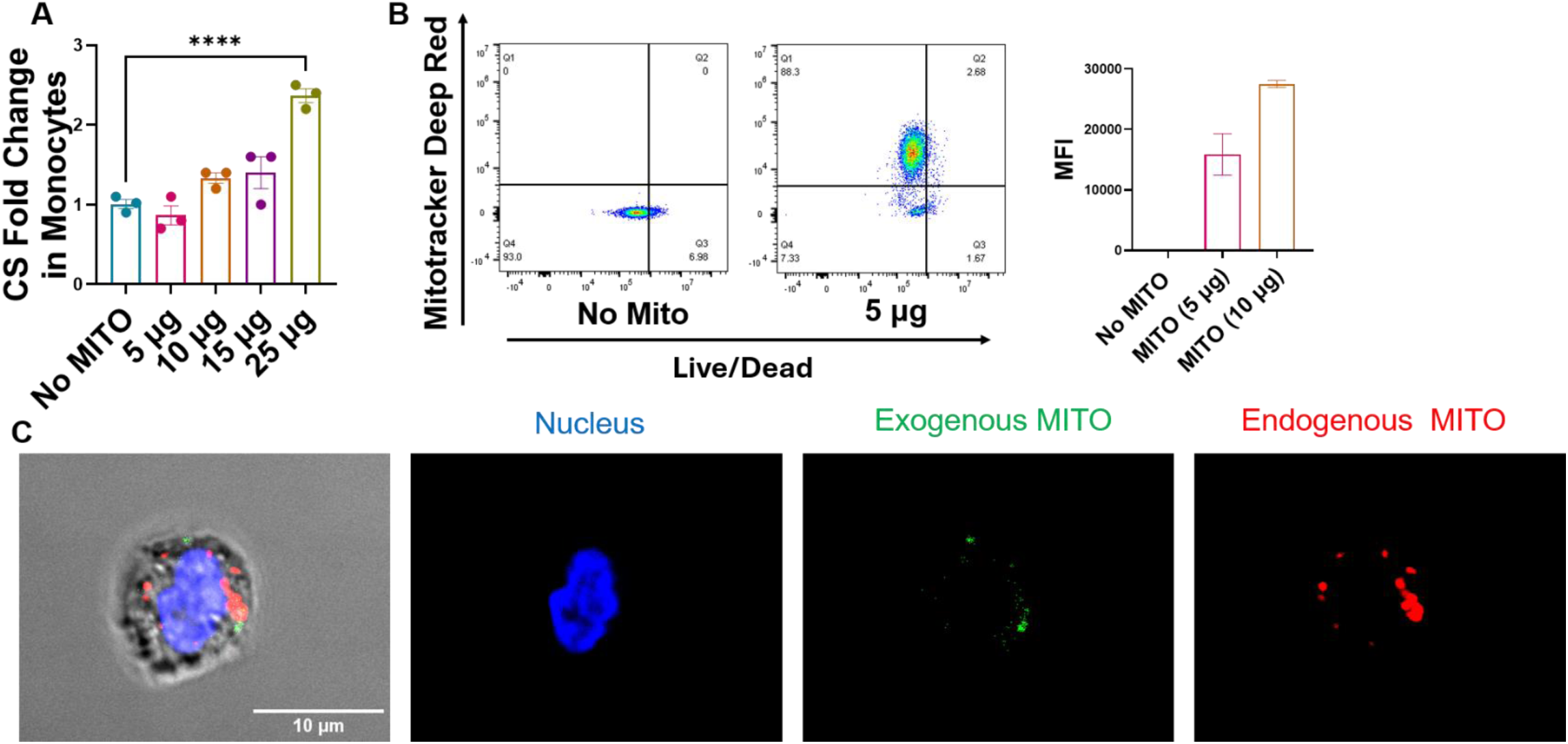
Human monocytes take up isolated mitochondria. (A) Citrate synthase activity in washed monocytes after 24 h of co-culture with isolated mitochondria at 5, 10, 15 and 25 µg of mitochondrial protein, expressed as fold change relative to untreated monocytes (n = 3). (B) Flow cytometric analysis of mitochondrial uptake and viability. Representative dot plots show MitoTracker Deep Red fluorescence (y axis) against viability dye (x axis), with quadrant gates set on untreated monocytes; the bar graph shows MitoTracker Deep Red MdFI in viable monocytes, which increased with the dose supplied (5 and 10 µg). (C) Representative confocal images of monocytes after 3 h of co-culture: merge with bright field showing the cell outline, nuclei (DAPI, blue), exogenous mitochondria (MitoTracker Green, green) and endogenous mitochondria (MitoTracker Deep Red, red), the latter labeled by pre-treating the monocytes before co-culture. Exogenous mitochondria lie within the monocyte boundary, close to the endogenous mitochondrial network. Scale bar, 10 µm. Data are mean ± SEM; (A) one-way ANOVA with Dunnett’s multiple comparisons test against untreated monocytes, ****P < 0.0001.

To measure uptake at single-cell resolution, we labeled isolated mitochondria with MitoTracker Deep Red before co-culture and analyzed the monocytes by flow cytometry. 88% of viable monocytes were MitoTracker Deep Red-positive after 3 h, and both the proportion of positive cells and the median fluorescence intensity increased with dose (0 to 50 µg; **Figure 2B** and **Supplementary Figure 2**), while viability was unchanged across the range tested.

To localize the transplanted organelles relative to the endogenous network, monocytes were pre-labeled with MitoTracker Deep Red to mark their own mitochondria and co-cultured for 3 h with isolated mitochondria (5 µg) labeled with MitoTracker Green, which accumulates in mitochondria largely independently of membrane potential. Confocal imaging showed exogenous mitochondria (green) within the monocyte boundary, lying close to the endogenous network (red) (**Figure 2C**). Imaging was performed at 3 h; functional readouts used 24 h of co-culture throughout. Monocytes carrying transplanted mitochondria are hereafter referred to as mitochondria-transplanted monocytes (MTM).

### Mitochondrial transplantation preserves viability and does not induce the measured basal inflammatory readouts

We next asked whether transplantation itself was toxic or activating, since isolated mitochondria carry damage-associated molecular patterns. Viability was assessed by annexin V and propidium iodide staining after 24 h of co-culture. Transplantation did not reduce the proportion of live monocytes, and the proportion of annexin V-positive cells was lower in MTM than in untreated monocytes (P = 0.0274, n = 3; **Supplementary Figure 3A, B**). Reduced annexin V staining indicates less phosphatidylserine exposure under these conditions and does not establish prolonged survival in vivo. Additionally, metabolic activity, measured by CCK-8, was maintained at 5 and 10 µg and fell at the higher dose tested (15 µg; P = 0.0342, n = 4; Supplementary Figure 3C).

To test whether the transplanted organelles triggered an inflammatory response, we assayed supernatants from recipient monocytes for TNF-α and IL-10 after 24 h. Neither cytokine showed a detectable increase above the level of untreated monocytes (n = 2 donors; **Supplementary Figure 4A, B**). We then used THP1-Dual reporter cells to interrogate NF-κB and IRF, the two pathways most likely to sense mitochondrial DAMPs. Mitochondria at 5 and 10 µg induced neither reporter, whereas Pam3CSK4 and LPS strongly induced NF-κB (17.6- and 4-fold; P < 0.0001 and P = 0.0009) and Pam3CSK4 induced IRF (1.9-fold; P = 0.010) (n = 4 independent experiments; **Supplementary Figure 4C, D**).

### Mitochondrial transplantation shifts monocytes toward a classical phenotype

To assess how transplantation alters monocyte phenotype and subset distribution, we immunostained untreated monocytes and MTM (5 µg of mitochondrial protein) after 24 h of co-culture and analyzed them by flow cytometry. Following the gating strategy shown in **Supplementary Figure 5**, subsets were defined on CD14 against CD16 as classical (CD14++CD16−), intermediate (CD14++CD16+), and non-classical (CD14+CD16++) (**Figure 3A**). Non-classical monocytes were rare, as expected after CD14-positive selection.

**Figure 3.**
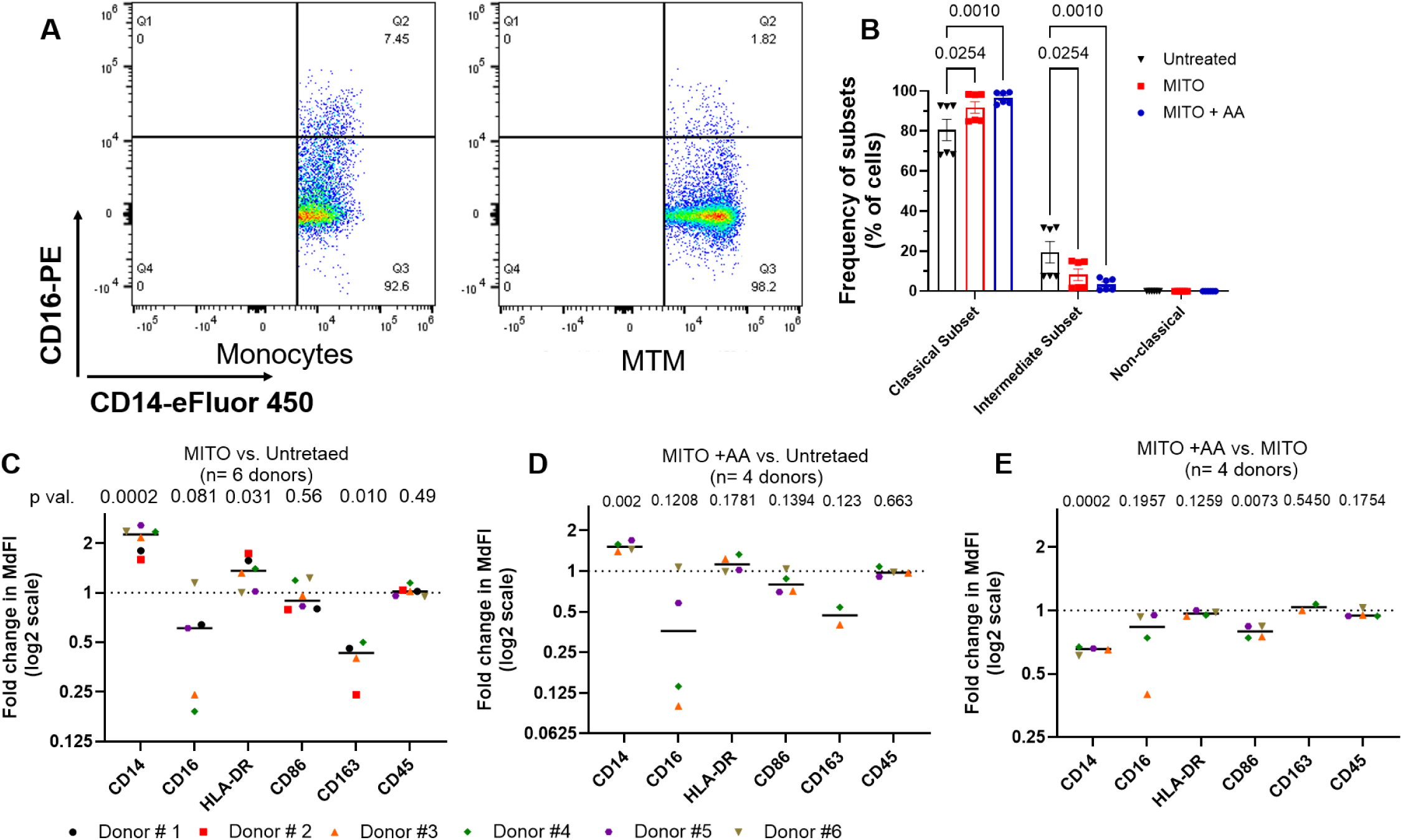
Mitochondrial transplantation shifts monocyte subset distribution and surface marker expression. Monocytes from six donors were cultured for 24 h alone, with 5 µg of isolated mitochondria (MTM), or with antimycin A-pretreated mitochondria (MTM+AA; donors 3–6), then analyzed by flow cytometry. (A) Representative dot plots of CD14 against CD16 in single, live monocytes, showing the classical (CD14++CD16−), intermediate (CD14++CD16+) and non-classical (CD14+CD16++) subsets. Quadrants were set on fluorescence-minus-one controls (Supplementary Figure 5). Non-classical monocytes were rare, as expected after CD14-positive selection. (B) Subset frequencies from the CD14/CD16 quadrants in six donors: the classical subset increased, and the intermediate subset decreased after transplantation. (C) MdFI of CD14, CD16, HLA-DR, CD86, CD163 and CD45 in MTM relative to matched untreated monocytes. CD14 rose 2.11-fold (95% CI 1.74–2.56, P = 0.0002; higher in 6 of 6 donors) and HLA-DR 1.31-fold (95% CI 1.04–1.66, P = 0.031); CD163 fell to 0.39-fold (95% CI 0.23–0.65, P = 0.010; lower in 4 of 4 donors in whom it could be quantified); CD16 fell to 0.46-fold without reaching significance (95% CI 0.18–1.16, P = 0.081; lower in 4 of 5); CD86 (0.95-fold, P = 0.56) and the CD45 staining control (1.02-fold, P = 0.49) were unchanged. (D) The same markers for MTM+AA relative to untreated monocytes (n = 4). CD14 rose 1.53-fold (95% CI 1.33–1.75, P = 0.0022) and no other marker changed significantly (P ≥ 0.12). (E) MTM relative to MTM+AA in the same four donors. CD14 (1.55-fold, P = 0.0002) and CD86 (1.27-fold, P = 0.0073) were higher with respiration-competent mitochondria, whereas CD16, HLA-DR, CD163 and CD45 did not differ (P ≥ 0.125). (B) Bars are mean ± SEM; two-way ANOVA with donor as a matching factor, followed by Tukey’s multiple comparisons test. (C–E) Each point is one donor, the horizontal bar is the mean of the donor fold changes and the dotted line at 1 marks no change; ratio paired t-tests on the donor-matched MdFI fold changes. P values shown are unadjusted; effect sizes with 95% confidence intervals and Benjamini–Hochberg q values across the six markers within each comparison are given in Supplementary Figure 6. Fold changes use only donors whose condition means remained positive after compensation (CD16, 5 donors in C; CD163, 4 in C and 2 in D and E).

Transplantation altered the subset distribution: the intermediate subset decreased, and the classical subset increased (P = 0.0254; **Figure 3B**, n = 6). Consistent with this, CD14 expression rose 2.11-fold relative to matched untreated monocytes (P = 0.0002, n = 6 donors), and CD16 fell to 0.46-fold, a decrease seen in four of five quantifiable donors but not significant (P = 0.081). HLA-DR rose 1.31-fold (P = 0.031) and CD163 fell to 0.39-fold (P = 0.010), while CD86 (0.95-fold, P = 0.56) and the CD45 staining control (1.02-fold, P = 0.49) did not differ significantly (**Figure 3C**). Supplementary Figure 6 summarizes all three comparisons with Benjamini–Hochberg-adjusted q values; after adjustment, CD14 and CD163 remain significant in this comparison.

To ask whether these changes require mitochondrial respiration, isolated mitochondria were treated with antimycin A for 30 min at room temperature and washed before co-culture (MTM+AA). Antimycin A-treated mitochondria still raised CD14 to 1.53-fold of untreated monocytes (P = 0.0022, n = 4 donors; **Figure 3D**), accounting for 49% of the total CD14 response. Direct comparison confirmed that CD14 (1.55-fold, P = 0.0002) and CD86 (1.27-fold, P = 0.008) were higher with respiration-competent mitochondria, whereas CD16, HLA-DR, CD163 and CD45 did not differ between the two (P ≥ 0.125; **Figure 3E**). Antimycin A is lipophilic and may have carried into the recipient cells despite washing, so this comparison does not partition the CD14 response between donor and host respiration.

### Mitochondrial transplantation reduces ATP-linked respiration with a trend toward greater relative respiratory reserve

We next asked whether transplantation alters recipient-cell metabolism. We measured oxygen consumption rate (OCR) and extracellular acidification rate (ECAR) in untreated monocytes, MTM, and MTM+AA using the Seahorse XF Cell Mito Stress Test, with sequential injection of oligomycin, FCCP, and rotenone/antimycin A (**Figure 4A, B**). Rates were normalized to post-assay Hoechst nuclear counts and expressed per 1,000 cells; each donor was assayed on two plates and contributes a single value (n = 4 donors).

**Figure 4.**
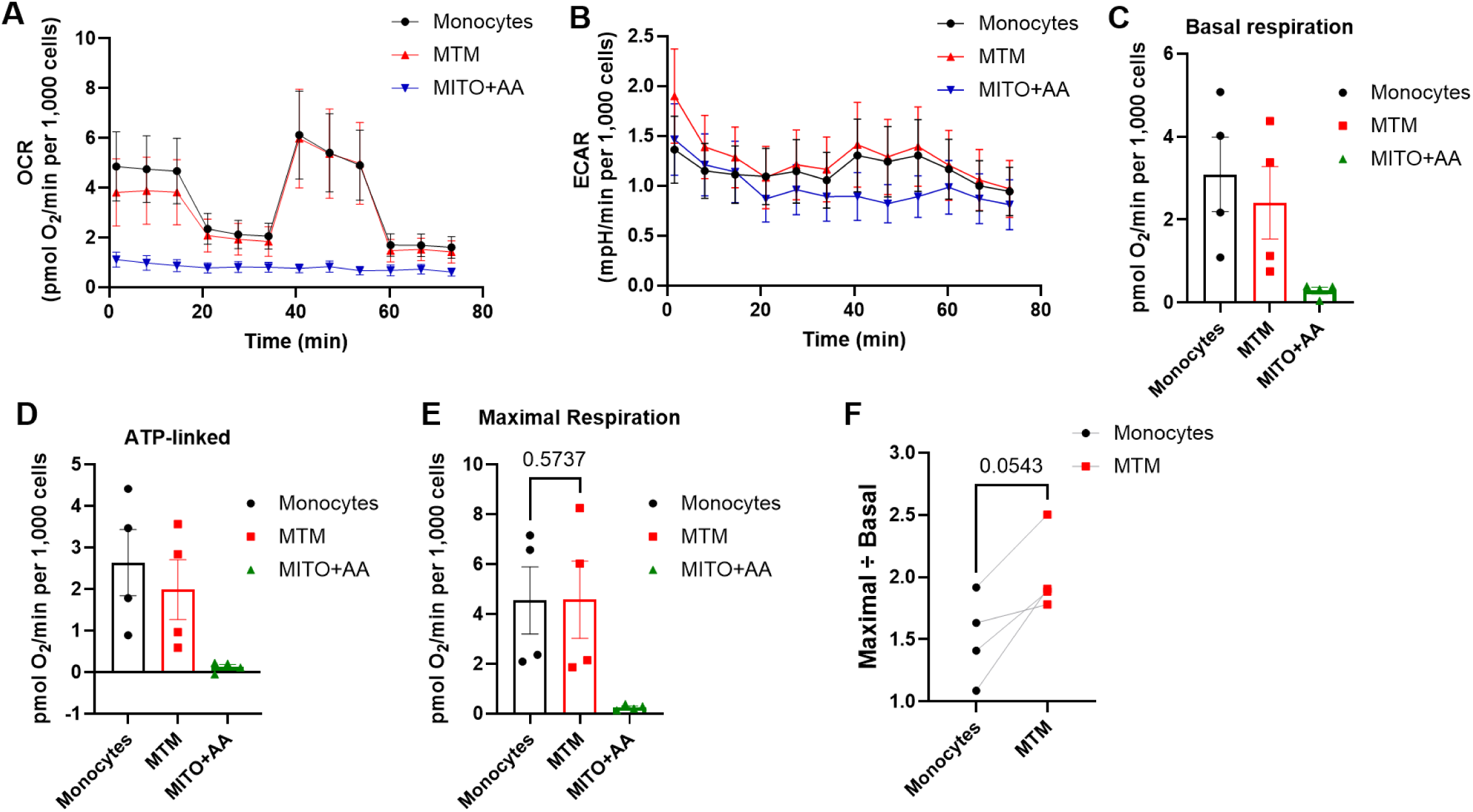
Mitochondrial transplantation reduces ATP-linked respiration with a trend toward greater relative respiratory reserve of human monocytes. Monocytes from four donors were cultured for 24 h in medium alone or with 5 µg of isolated mitochondria (MTM) or antimycin A-pretreated mitochondria (MTM+AA), then assayed with the Seahorse XF Cell Mito Stress Test. (A) Oxygen consumption rate (OCR) and (B) extracellular acidification rate (ECAR) over time; lines mark the injection of oligomycin, FCCP and rotenone/antimycin A. (C) Basal respiration, (D) ATP-linked respiration, (E) maximal respiration and (F) relative respiratory reserve (maximal ÷ basal); MTM+AA is omitted from (F) because respiration was abolished in that group. Rates are normalized to post-assay Hoechst nuclear counts and expressed per 1,000 cells. Each point is one donor (n = 4) and lines join the same donor; bars are means; (A, B) show mean ± SEM. P values are from ratio paired t-tests on donor means and are unadjusted.

Transplantation reduced ATP-linked respiration, with a nonsignificant reduction in basal respiration. Basal respiration in MTM was 0.72-fold of matched untreated monocytes (P = 0.0543) and ATP-linked respiration 0.72-fold (P = 0.044), while maximal respiration did not differ significantly (0.97-fold, P = 0.61) (**Figure 4C–E**). Expressed as a ratio, maximal divided by basal respiration rose 1.34-fold (P = 0.0543); expressed in absolute terms, spare respiratory capacity, maximal minus basal respiration, rose by 0.72 pmol O₂/min per 1,000 cells (P = 0.17). Neither reached significance, but both were higher in all four donors (**Figure 4F**). Monocytes given antimycin A-treated mitochondria showed almost no respiration: basal 0.10-fold, ATP-linked 0.03-fold, and maximal 0.06-fold of untreated monocytes (all P ≤ 0.017), and FCCP produced no stimulation above baseline.

To ask whether the metabolic change persists, monocytes and MTM were assayed after 24 h and after 6 days in culture (five independent experiments per arm; **Supplementary Figure 7**). These plates were not normalized to cell number, so comparisons were restricted to within-well ratios, which do not account for changes in viability or cell composition. The respiratory control ratio, maximal divided by proton leak, rose at 24 h (1.74-fold, P = 0.011) and fell after 6 days (0.28-fold, P = 0.030), with proton leak 15.7-fold higher than control (P = 0.005) and FCCP failing to raise OCR above baseline in four of five MTM wells. Maximal divided by basal respiration showed the same reversal (0.66-fold at 6 days, P = 0.060). The separate extended-culture experiments indicate that the respiratory phenotype changes over time and does not establish a sustained benefit.

### Mitochondrial transplantation redirects the monocyte response to LPS

We next asked whether the phenotypic and metabolic changes were accompanied by a change in inflammatory function. We challenged monocytes and MTM with LPS (10^3^ EU/ml) for 24 h and measured 13 cytokines and chemokines in the supernatant using a BioLegend LEGENDplex bead-based immunoassay (n = 6 donors). Results are expressed relative to each donor’s LPS-stimulated untreated monocytes.

Transplantation markedly suppressed TNF-α. Release fell to 0.06-fold of untreated monocytes (one-tailed P < 0.0001) and was lower in all six donors (**Figure 5A**). CXCL10/IP-10 was also reduced to 0.25-fold (P = 0.0241; lower in 5 of 6 donors) (**Figure 5B**). Conversely, IL-10 rose 3.33-fold (higher in 5 of 6 donors; one-tailed P = 0.0404) (**Figure 5C**). The IL-10 response varied considerably between donors (0.65-to 34-fold); the one donor whose IL-10 did not rise showed normal suppression of TNF-α and IP-10, indicating a selective rather than a global failure to respond. IL-2 (0.96-fold, P = 0.634; **Figure 5D**) and IL-6 (1.10-fold, P = 0.869; **Figure 5E**) did not differ significantly. The overall change in profile is summarized in **Figure 5F**. Among the remaining analytes, MCP-1/CCL2 showed a modest and directionally consistent increase (1.20-fold, higher in 5 of 6 donors) that did not reach significance (P = 0.10), and no other cytokine or chemokine showed a statistically significant difference between transplanted and untreated monocytes (P ≥ 0.36, **Supplementary Figure 8**). The IL-10 increase did not reach significance under two-tailed testing. IL-1β, IL-17A, and TGF-β1 were below the limit of quantification in five of six donors, precluding reliable conclusions about these analytes.

**Figure 5.**
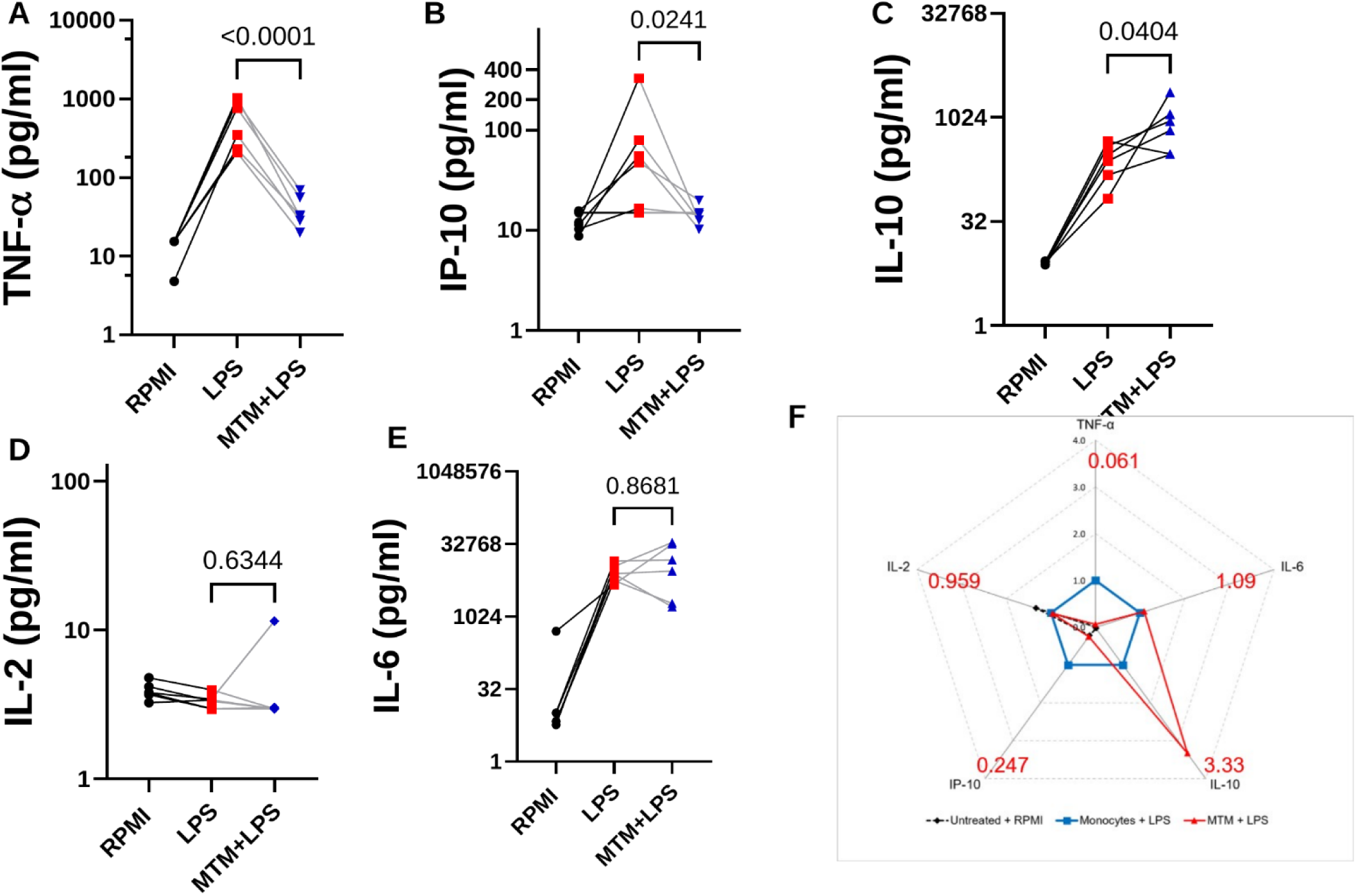
Mitochondrial transplantation redirects the inflammatory response of human monocytes to LPS. Monocytes were cultured in RPMI alone, stimulated with LPS (10³ EU/ml), or given 5 µg of isolated mitochondria for 24 h and then stimulated with LPS (mitochondria-transplanted monocytes, MTM+LPS). Supernatants collected after 24 h of LPS stimulation were assayed for 13 cytokines and chemokines with a BioLegend LEGENDplex bead-based immunoassay. (A) TNF-α, (B) CXCL10/IP-10, (C) IL-10, (D) IL-2 and (E) IL-6 in six independent donors. These five were selected on biological grounds as the analytes most relevant to monocyte activation; the remaining eight are shown in Supplementary Figure 8. (F) Profile of cytokine release across the same five analytes, summarized as fold change relative to LPS-stimulated untreated monocytes (linear radial axis). Values at each vertex are geometric mean fold changes across donors: TNF-α 0.061, IP-10 0.247, IL-10 3.33, IL-6 1.10 and IL-2 0.959. Each point is one donor and lines join the same donor across conditions; technical replicates were averaged within donor, so n = 6 donors throughout. P values compare MTM+LPS with LPS alone by ratio paired t-test on donor means; TNF-α and IL-10 were tested one-tailed under a direction specified before analysis, and all other analytes two-tailed.

Mitochondria alone did not induce TNF-α or IL-10 (**Supplementary Figure 4A, B**), so these changes reflect a shift in how the cells respond to stimulation rather than a response to the transplant itself. In the three donors assayed with both arms, monocytes given antimycin A-treated mitochondria tended to release more TNF-α than those given respiration-competent mitochondria (0.53-fold for MTM relative to MTM+AA, P = 0.21), while IL-10 did not differ (1.09-fold, P = 0.66) (**Supplementary Figure 8I**). Phagocytosis, an energy-dependent effector function, was unchanged: uptake of pHrodo-labeled E. coli particles did not differ between MTM and untreated monocytes (0.86-fold, P = 0.68, n = 5 donors; Supplementary Figure 9A), a readout that reflects both particle internalization and phagosomal acidification. Rotenone pre-treatment reduced uptake in both groups, and in the four donors assayed with both preparations uptake was 1.27-fold higher with respiration-competent than with antimycin A-treated mitochondria (P = 0.011; Supplementary Figure 9B). Together, these results show that mitochondrial transplantation reshapes rather than silences the monocyte response to LPS: TNF-α and IP-10 release is strongly suppressed, IL-10 increases in most donors, and monocyte chemoattractant output is preserved or modestly raised.

## Discussion

In this study, transplantation of freshly isolated, respiration-competent mitochondria altered the surface phenotype, respiratory profile and LPS responsiveness of primary human monocytes. The resulting mitochondria-transplanted monocytes (MTM) showed a reduced frequency of the intermediate subset, with increased CD14 and HLA-DR and decreased CD163; reduced ATP-linked respiration with maximal capacity preserved; and, after LPS stimulation, markedly less TNF-α and CXCL10/IP-10 with more IL-10 in most donors. Together these findings show that mitochondrial transplantation modifies monocyte subsets and selectively alters inflammatory mediator production.

Each altered marker has a defined function, and the literature on them limits what the changes can show. CD14 facilitates LPS recognition by the TLR4–MD-2 complex,^27^ HLA-DR participates in MHC class II-mediated antigen presentation,^28^ and CD163 is the hemoglobin–haptoglobin scavenger receptor, whose surface abundance is also influenced by proteolytic shedding.^29^ In human intestinal macrophages, high CD14 with low CD163 is characteristic of the inflamed rather than the homeostatic state,^7,30^ and CD14⁺CD163ˡᵒʷ cells isolated from Crohn’s disease mucosa show increased Th17-inducing activity;^9^ on those two markers alone MTM would resemble a population associated with intestinal inflammation, yet their functional output runs in the opposite direction. The same caution applies to the macrophage polarization framework, which was defined in cells differentiated over six to eight days^31,32^ rather than at 24 h. IL-10-driven polarization (M2c) combines increased CD14 and CD163 with high IL-10 and low inflammatory cytokine output;^33^ MTM share three of those features but not the fourth, since CD163 fell rather than rose. In our own data the clearest change in subset distribution was a reduced frequency of intermediate monocytes alongside lower CD16 expression. The CD16 decrease was large (0.46-fold); because the classical and intermediate gates are separated on the CD16 axis, the two measures are not independent, and the subset shift should be read as a consequence of lower CD16 rather than as conversion to a stable classical identity, while CD14-positive selection restricted assessment of non-classical monocytes and limits generalization to the complete circulating pool. These changes document remodeling of recognition and antigen-presentation machinery without implying functional enhancement, and rather than forcing MTM into an existing category we would argue that the functional readouts define this state, with the divergence at CD163 a feature worth characterizing in its own right.

The respiratory profile suggests a shift in the balance between ongoing respiratory activity and available reserve. MTM showed reduced ATP-linked respiration and a trend toward greater relative respiratory reserve. The pattern is therefore consistent with greater reserve relative to demand rather than with a raised respiratory ceiling. That distinction matters for monocyte adaptation: in glucose-deprived, LPS-stimulated human monocytes, basal and ATP-linked respiration rose while maximal respiration and spare capacity fell,^34^ showing how inflammatory demand draws on available capacity. Because classical and non-classical monocytes also differ in their metabolic programs,^35^ and because measurements of the CD14-selected population cannot distinguish changes within individual subsets from effects of their relative abundance, the respiratory findings should be read alongside the altered subset frequencies. Antimycin A treatment significantly reduced donor-organelle membrane potential, yet monocytes receiving these preparations lost respiration almost entirely, exceeding what the transplanted organelles alone could account for; carryover of inhibitor onto the recipient respiratory chain and uptake of damaged organelles could both contribute, and residual inhibitor was not assayed. The comparison therefore shows that respiration-competent and respiration-incompetent preparations give different outcomes, without attributing the difference to respiration of the donor mitochondria. The organelle source is a further untested variable: HEK293T cells were chosen for yield and consistency, but they are a transformed line whose mitochondria may differ from those of primary cells in respiratory capacity per unit protein, mitochondrial DNA copy number, membrane composition and haplotype. The fate of the transplanted organelles and their integration with the endogenous network also remain unresolved.

The functional change was selective rather than global. TNF-α release was markedly reduced in all six donors and CXCL10/IP-10 in five, IL-10 increased in five with considerable variation in magnitude, and most remaining mediators showed no detectable difference. Suppression of TNF-α and IP-10 also occurred in the donor whose IL-10 did not increase, so these effects do not require a detectable rise in endpoint IL-10, although this does not exclude a role for basal or earlier IL-10 signaling. The donor cohorts are modest, no multiplicity correction was applied across the panel. One feature of this result requires comment. CD14 is the co-receptor that transfers LPS to the TLR4–MD-2 complex, and it is also required for LPS-induced TLR4 endocytosis and the TRIF-dependent arm of signaling that generates CXCL10/IP-10.^36,37^ MTM expressed more than twice as much CD14 as matched untreated monocytes, yet released markedly less of both the principal MyD88-dependent output and the principal TRIF-dependent one.

Several explanations are possible and are not mutually exclusive. Altered subset composition may contribute, since intermediate monocytes are a principal source of TNF-α^38^ and their frequency fell, but the magnitude of the cytokine change exceeds what the loss of a minority subset can readily explain. Alternatively, the additional CD14 may be a consequence of an anti-inflammatory program rather than a cause of altered LPS sensitivity, since IL-10 and related conditioning signals induce CD14 on human monocytes while suppressing TNF-α production;^33^ that reading is also incomplete, because such conditioning characteristically raises CD163, which fell here. An autocrine IL-10 loop would be the most direct explanation, but suppression was equally evident in the donor whose IL-10 did not rise. Distinguishing these possibilities will require determining whether the additional CD14 is functionally competent, for instance by measuring LPS binding or TLR4 internalization. Both suppressed mediators are implicated in intestinal inflammation, where the efficacy of anti-TNF antibodies depends on IL-10-dependent regulatory macrophage differentiation^39^ and CXCL10 production is elevated in active disease.^40,41^ The reduced intermediate frequency is of related interest, since that subset is expanded in active Crohn’s disease^10^ and CD14⁺CD16⁺ monocyte-derived populations accumulate in inflamed mucosa.^8^ Because maturation of recruited monocytes into relatively TLR-unresponsive resident macrophages is arrested during experimental colitis,^6^ and circulating monocytes from patients with Crohn’s disease already carry inflammatory programming,^11^ conditioning cells before recruitment is worth testing, although that would require establishing that the MTM phenotype persists through differentiation. Validation in monocytes from patients with IBD, and evaluation of alternative mitochondrial donor sources, will be needed to establish disease relevance and generalizability.

### Conclusions

This study demonstrates that transplantation of functional mitochondria modifies the surface phenotype, respiratory profile, and inflammatory responsiveness of primary human monocytes. Reduced TNF-α and CXCL10/IP-10 release following LPS stimulation, accompanied by increased IL-10 in most donors, identifies a selective rather than a global change in inflammatory output. These findings identify the circulating monocyte as a tractable target for mitochondria-based modulation of inflammation, and provide a basis for investigating whether such conditioning influences the behavior of monocyte-derived macrophages in tissue, including in the intestine.

## Materials and Methods

Methods for materials and reagents, confocal imaging, flow cytometry and surface immunostaining, flow cytometric uptake, Seahorse analysis of monocyte respiration, cell death, metabolic activity, THP1-Dual reporter assays and phagocytosis, together with full statistical detail, are provided in the Supporting Information.

### Human donors and experimental design

De-identified whole blood from healthy adult donors was purchased from Research Blood Components LLC (Boston, MA, USA), which collects under its own institutional review board-approved protocol with written informed consent from all donors. Because the material was commercially obtained and fully de-identified, with no access to identifiable private information, this work did not constitute human subjects research and did not require additional institutional review. Blood was collected into K2EDTA. Primary monocyte experiments used a donor-matched design, with the same donor contributing cells to the treatment conditions compared within each experiment. The number of donors varied by assay and is specified in the corresponding figure legends.

### Mitochondrial isolation and characterization

We isolated mitochondria from HEK293T cells using an established mechanical homogenization and differential centrifugation procedure.^23,42,43^ HEK293T cells were cultured as described above. At approximately 80% confluence, cells were detached using 1% trypsin, collected by centrifugation, and washed once with ice-cold 1× phosphate-buffered saline. We resuspended cell pellets in mitochondrial isolation buffer containing 10 mM HEPES (pH 7.5), 25 mM sucrose, 1 mM ATP, 0.1 mM ADP, 5 mM sodium succinate, and 2 mM K₂HPO₄. We mechanically disrupted cell suspensions using a pestle-driven homogenizer, then centrifuged the homogenate at 700 × g for 5 min to remove cellular debris. The supernatant was collected and centrifuged at 8,000 × g for 5 min. The resulting pellet was resuspended in the same isolation buffer and centrifuged again at 8,000 × g for 10 min. The final mitochondria-enriched pellet was collected for subsequent characterization and transplantation. Protein concentration was determined using the Pierce Bradford Plus Protein Assay Kit (Thermo Fisher Scientific; Cat. No. A55866), and citrate synthase activity was measured using the Citrate Synthase Assay Kit (Sigma-Aldrich; Cat. No. CS0720), according to the manufacturers’ instructions.

### Seahorse analysis of isolated mitochondria

We assessed respiratory function in freshly isolated mitochondria using a Seahorse XF HS Mini analyzer (Agilent Technologies), following a protocol adapted from Yang et al.^44^ Mitochondrial assay solution (MAS) consisted of 220 mM D-mannitol, 70 mM sucrose, 10 mM KH2PO4, 5 mM MgCl2, 1 mM EGTA, and 2 mM HEPES. The buffer was adjusted to pH 7.2 with 1 M KOH and filter-sterilized (0.22 µm) before experimental use with bovine serum albumin (BSA, 0.2% (w/v)), ADP, pyruvate, and malate. The MAS/BSA premix contained 0.4% (w/v) BSA before addition of substrate stocks, and the assay targets were 4 mM ADP, 5 mM pyruvate, and 0.5 mM malate. Mitochondria were loaded at 4 µg of mitochondrial protein in 25 µL per well, and the plate was centrifuged at 2,000 × g for 20 min at 4 °C. Subsequently, 155 µL of assay medium was added to each well, bringing the initial assay volume to 180 µL. The plate was equilibrated for 5 min at 37 °C before analysis. Oxygen consumption rate (OCR) and extracellular acidification rate (ECAR) were recorded using cycles comprising 90 s mixing, no waiting period, and 120 s measurement. Following initial measurements, oligomycin, FCCP, and rotenone/antimycin A were injected sequentially at nominal final well concentrations of 3.16 µM oligomycin, 4 µM FCCP, 4 µM antimycin A, and 1 µM rotenone. The FCCP injectate additionally contained 1 mM pyruvate. We used medium-only wells for background correction, and normalized rates to mitochondrial protein content.

Because ADP was present from the start of the assay, the initial OCR represented ADP-stimulated respiration. State 3 respiration was calculated as initial OCR minus residual OCR after rotenone/antimycin A addition. ATP-linked respiration was calculated as the difference between initial and oligomycin-inhibited OCR. Leak respiration was calculated as oligomycin-insensitive OCR minus residual OCR following rotenone/antimycin A addition. Maximal uncoupled respiration (State 3u) was calculated as peak FCCP-stimulated OCR minus residual OCR. Coupling efficiency was calculated as ATP-linked respiration divided by residual-corrected State 3 respiration, multiplied by 100. Respiratory control ratios were calculated as State 3u/leak and, separately, State 3/leak. The difference between State 3u and State 3 respiration was reported as the respiratory margin between ADP-stimulated and uncoupled conditions.

### Assessment of mitochondrial membrane potential

Mitochondrial membrane potential was assessed using JC-1 (Molecular Probes, Thermo Fisher Scientific; Cat. No. T3168). Aliquots containing 10 or 50 µg of isolated mitochondrial protein were incubated with JC-1 at a final concentration of 2 µM for 30 min at room temperature. Fluorescence was measured using a SpectraMax® iD3 plate reader, with excitation at 488 nm and emission at 525 nm (green) and 590 nm (red). Membrane potential was evaluated using the red-to-green fluorescence ratio (F₅₉₀/F_520_). Untreated mitochondria were compared with antimycin A-treated preparations (1 µM for 30 min at room temperature, followed by two washes in MITO buffer with centrifugation at 8,000 × g for 10 min).

### Monocyte isolation and mitochondrial transplantation

Peripheral blood mononuclear cells (PBMCs) were isolated by density-gradient centrifugation using Lymphoprep (STEMCELL Technologies), according to the manufacturer’s instructions. CD14⁺ monocytes were subsequently isolated by positive immunomagnetic selection using the EasySep™ Human CD14 Positive Selection Kit II (STEMCELL Technologies; Cat. No. 17858), following the manufacturer’s protocol. Isolated monocytes were seeded in 96-well culture plates at 1 × 10⁵ cells per well and incubated for 30 min at 37 °C in a CO₂ incubator before mitochondrial addition unless otherwise noted. We added freshly isolated mitochondria at 5, 10, or 15 µg of mitochondrial protein per well. We co-incubated monocytes and mitochondria for 24 h under the same culture conditions. Donor-matched monocytes cultured without added mitochondria served as untreated controls. Monocytes receiving isolated mitochondria were designated mitochondria-transplanted monocytes (MTM).

To assess mitochondrial content after transplantation, we measured citrate synthase activity in washed monocytes (10^5^ cells per well in 96-well plate) after 24 h of co-culture using the Citrate Synthase Assay Kit (Sigma-Aldrich; Cat. No. CS0720), the same kit used to characterize the isolated mitochondrial preparations. We performed assays according to the manufacturer’s instructions and compared activity with that of donor-matched untreated monocytes.

### Cytokine and chemokine measurements

Cytokine measurements used donor-matched monocytes cultured with or without mitochondria, with unstimulated controls analyzed in parallel. Following 24 h of co-culture with mitochondria, the supernatant was decanted and replaced with fresh medium alone or fresh medium containing LPS (1 × 10³ EU/mL), to reduce extracellular mitochondrial carryover before stimulation. Cells were incubated for a further 24 h at 37 °C. Supernatants were then collected and stored at −20°C until analysis. Supernatants were also collected in parallel from unstimulated monocytes cultured with or without mitochondria, to assess cytokine release in the absence of LPS. Cytokine and chemokine concentrations were quantified using the LEGENDplex™ Human Essential Immune Response Panel (13-plex; BioLegend), according to the manufacturer’s instructions.^45^ Samples were acquired using an Attune CytPix flow cytometer (Thermo Fisher Scientific), and concentrations were determined from analyte-specific standard curves. Technical replicates were averaged within each donor and treatment condition before statistical analysis.

## Statistical analysis

Data were analyzed using GraphPad Prism 11. The biological replicate was an individual donor for primary monocyte experiments, an independent mitochondrial preparation for isolated mitochondrial assays, and an independent experiment for THP1-Dual reporter assays; technical replicates were averaged within each biological replicate before analysis, and donor matching was retained in comparisons between treatment conditions. Donor-matched fold changes were compared by ratio paired t-tests on log-transformed values, with geometric means and 95% confidence intervals back-transformed for presentation, and a Benjamini–Hochberg adjustment was applied across the six surface markers within each comparison. Tests were two-tailed except for TNF-α and IL-10 in the LPS response, for which a direction was specified before analysis; no correction was made across the 13 analytes. A P value below 0.05 was considered statistically significant.

## Author Contributions

Saeed Nazemidashtarjandi (Conceptualization, Methodology, Formal analysis, Investigation, Data curation, Writing—Original draft, Writing—Review & Editing, Supervision, Visualization, Funding acquisition), Sairam Pantham (Formal analysis, Writing— Review & Editing), Rachel Dori (Formal analysis, Writing—Review & Editing), Martin Yarmush (Conceptualization, Writing—Review & Editing, Supervision, Funding acquisition).

## Data Availability Statement

The data that support the findings of this study are available in the Materials and Methods, Results, and/or Supplemental Material of this article.

