## Supporting Information for "Mitochondrial transplantation redirects rather than suppresses the inflammatory response in primary human monocytes"

**Supplementary Materials and Methods**

**Materials**

Ultrapure lipopolysaccharide from *Escherichia coli* (LPS-EB Ultrapure; InvivoGen, San Diego, CA, USA; Cat. No. tlrl-3pelps) was reconstituted at $5 \times 10^6$ endotoxin units (EU)/mL. Pam3CSK4, a synthetic TLR1/2 agonist, was obtained from InvivoGen (Cat. No. tlrl-pms). Primary human monocytes were cultured in RPMI 1640 medium (Gibco, Thermo Fisher Scientific; Cat. No. 11875119; purchased from Fisher Scientific) supplemented with 10 μg/mL gentamicin (Sigma-Aldrich; Cat. No. G1397-10ML), 10 mM pyruvate, 10 mM HEPES, and 10% (v/v) pooled human AB serum. The serum was derived from male AB clotted whole blood, was of USA origin, and was sterile-filtered (Sigma-Aldrich; Cat. No. H6914-20ML). Glucose (5.5 mM) used for culture supplementation was obtained from Sigma-Aldrich. THP1-Dual™ reporter cells were obtained from InvivoGen (Cat. No. thpd-nfis) and maintained according to the manufacturer's recommended culture protocol.

**Reagents and assays**

Peripheral blood mononuclear cells were isolated using Lymphoprep™ (STEMCELL Technologies, Vancouver, BC, Canada, Cat. No. 07861). We then isolated human $CD14^+$ monocytes by positive immunomagnetic selection using the EasySep™ Human CD14 Positive Selection Kit II (STEMCELL Technologies; Cat. No. 17858). Fluorophore-conjugated anti-human monoclonal antibodies were obtained from Thermo Fisher Scientific (Waltham, MA, USA) and included PE-conjugated anti-CD16 (clone CB16; Cat. No. 12-0168-42), eFluor 450-conjugated anti-CD14 (clone 61D3; Cat. No. 48-0149-42), APC-conjugated anti-CD45 (clone HI30; Cat. No. 17-0459-42), eFluor 506-conjugated anti-HLA-DR (clone LN3; Cat. No. 69-9956-42), Alexa Fluor 488-conjugated anti-CD86 (clone IT2.2; Cat. No. 53-0869-42), and PE-Cyanine7-conjugated anti-CD163 (clone eBioGHI/61; Cat. No. 25-1639-42). Cell Staining Buffer was obtained from

BioLegend (San Diego, CA, USA; Cat. No. 420201). We identified nonviable cells using eBioscience™ Fixable Viability Dye eFluor 780 (Thermo Fisher Scientific; Cat. No. 65-0865-14).

We determined mitochondrial protein concentration using the Pierce™ Bradford Plus Protein Assay Kit with Dilution-Free BSA Protein Standards (Thermo Fisher Scientific; Cat. No. A55866). We measured citrate synthase activity using the Citrate Synthase Assay Kit (Sigma-Aldrich; Cat. No. CS0720). MitoTracker™ Deep Red and MitoTracker™ Green probes (Thermo Fisher Scientific; Cat. No. M46753, M46750) were used for mitochondrial labeling. We assessed mitochondrial membrane potential using Molecular Probes™ JC-1 Dye (Thermo Fisher Scientific; Cat. No. T3168).

Cytokines and chemokines were quantified using the LEGENDplex™ Human Essential Immune Response Panel (13-plex; BioLegend, V-bottom plate; Cat. No. 740930).[1] QUANTI-Blue™ and QUANTI-Luc™ reagents (InvivoGen; Cat. No. rep-qbs2, rep-qlc4lg2) were used to detect NF-κB-inducible SEAP and IRF-inducible Lucia luciferase reporter activities, respectively. Phagocytosis-associated fluorescence was assessed using pHrodo™ Deep Red *E. coli* BioParticles™ Conjugate for Phagocytosis (Thermo Fisher Scientific; Cat. No. P35360). Hoechst 33342 (Thermo Fisher Scientific; Cat. No. H3570) was used for post-assay nuclear staining and cell counting. Respiratory assays used oligomycin, carbonyl cyanide *p*-trifluoromethoxyphenylhydrazone (FCCP), rotenone, and antimycin A.

HEK293T cells were obtained from the American Type Culture Collection (ATCC) and cultured in high-glucose Dulbecco's modified Eagle medium with L-glutamine and sodium pyruvate (HyClone; Cat. No. SH30243.01) supplemented with 10% fetal bovine serum (MedSupply Partners; Cat. No. 62-1300-2HI) and 1% penicillin–streptomycin (Invitrogen, Thermo Fisher Scientific; Cat. No. 15140122).

**Confocal imaging of mitochondrial uptake**

For confocal imaging, monocytes were pre-labeled with MitoTracker™ Deep Red (1 μM) for 30 min at 37 °C to visualize endogenous mitochondria and subsequently washed with PBS to remove unbound dye. Isolated mitochondria were labeled separately with MitoTracker™ Green (1 μM) for 30 min at 37 °C, followed by two washes in MITO buffer with centrifugation at 8,000 × g for 10 min. Labelled mitochondria were added to pre-labeled monocytes at 5 μg of mitochondrial protein, and co-incubated for 3 h at 37 °C. Cells were then fixed with formalin (4% v/v) for 10 min. Images were acquired using an Olympus FV1000-MPE confocal microscope equipped with a 60× water-immersion objective to examine the intracellular localization of exogenous mitochondria relative to the endogenous mitochondrial network.

**Flow cytometry and surface immunostaining**

Following 24 h of co-culture, monocytes from each treatment condition were stained with Fixable Viability Dye eFluor 780 (1:2,000) for 30 min on ice and washed to remove excess dye. Before surface immunostaining, cells were blocked for 20 min in phosphate-buffered saline containing bovine serum albumin to reduce nonspecific antibody binding. Cells were then stained for 30 min on ice in Cell Staining Buffer (BioLegend, San Diego, CA, USA) in a total volume of 200 μL per sample, using 5 μL of each antibody against CD45, CD14, CD16, CD86, CD163 and HLA-DR. Antibody clones, fluorophores and catalogue numbers are given in the Materials section. Compensation was set using UltraComp eBeads™ Spectral Unmixing Beads (Cat. No. U20250). Following staining, cells were washed to remove unbound antibody and analyzed on a four-laser Attune CytPix flow cytometer equipped with a plate loader (Thermo Fisher Scientific), acquiring 20,000 events per sample.

Data were analyzed using FlowJo version 10 (BD Biosciences). We excluded debris, doublets, and nonviable cells before identifying monocytes based on CD45, HLA-DR, CD14, and CD16 expression. Within the viable singlet monocyte population, we classified subsets as classical ($CD14^{++}CD16^{-}$), intermediate ($CD14^{++}CD16^{+}$), or non-classical ($CD14^{+}CD16^{++}$), as shown in

Supplementary Figure 5. We acquired fluorescence-minus-one (FMO) controls for CD14 and CD16 to support gate placement. Subset frequencies and surface-marker expression were assessed within the gated monocyte population, with marker expression summarized as median fluorescence intensity (MdFI).

**Flow cytometric assessment of mitochondrial uptake**

Isolated mitochondria were labeled with MitoTracker™ Deep Red at 1 μM for 30 min at 37 °C. Unbound dye was removed by washing in mitochondrial isolation buffer with centrifugation at 8,000 × g for 10 min before co-culture with primary human monocytes. Monocytes were exposed to labeled mitochondria for 3 h at 37 °C over a dose range of 0–50 μg of mitochondrial protein. Following incubation, recipient monocytes were washed by centrifugation at 300 × g for 10 min and analyzed using an Attune CytPix flow cytometer (Thermo Fisher Scientific).

Debris, doublets, and nonviable cells were excluded before analysis. The threshold for MitoTracker Deep Red positivity was established using untreated monocytes as the negative control. The percentage of Deep Red-positive cells was calculated within the viable singlet monocyte population, and median fluorescence intensity (MdFI) was quantified within the viable singlet monocyte population. These measurements assessed acquisition of exogenous mitochondrial fluorescence; intracellular localization was examined separately by confocal microscopy.

**Seahorse analysis of monocyte respiration**

We assessed cellular respiration using a Seahorse XF HS Mini analyzer (Agilent Technologies) with the Seahorse XF Cell Mito Stress Test. Monocytes from four independent donors were seeded directly into Seahorse culture miniplates and cultured for 24 h in medium alone, with isolated mitochondria (5 μg of mitochondrial protein; MTM), or with an equivalent dose of antimycin A-pretreated mitochondria (MTM+AA). For MTM+AA, isolated mitochondria were treated with antimycin A (1 μM) for 30 min at room temperature and then washed twice in mitochondrial isolation buffer, pelleting at 8,000 × g for 10 min between washes, before co-culture. Before the assay, plates were centrifuged at 300 × g for 5 min and the medium was replaced with Seahorse XF RPMI assay medium supplemented with 5.5 mM glucose, 2 mM L-glutamine and 1 mM sodium pyruvate; plates were then equilibrated for [duration] at 37 °C in a non-$CO_2$ incubator. Oxygen consumption rate (OCR) and extracellular acidification rate (ECAR) were recorded before and after sequential injection of oligomycin (1.5 μM), FCCP (1 μM), and rotenone (1 μM) with antimycin A (2 μM), using the mixing, waiting, and measurement intervals specified in the instrument protocol. Sodium pyruvate (1 mM) was co-injected with FCCP to support maximal respiration. For the cell-normalized experiment, background-corrected OCR and ECAR were normalized to post-assay Hoechst-labeled nuclear counts quantified using QuPath and expressed per 1,000 cells. Each donor was assayed on two plates, and replicate measurements were combined within donor and treatment condition so that each donor contributed a single value per condition.

Non-mitochondrial oxygen consumption was defined as residual OCR following rotenone/antimycin A addition. Basal mitochondrial respiration was calculated as pre-injection OCR minus residual OCR. ATP-linked respiration was calculated as the oligomycin-sensitive component of OCR, and proton leak as oligomycin-insensitive OCR minus residual OCR. Maximal respiration was calculated from FCCP-stimulated OCR after subtraction of residual OCR. Relative respiratory reserve was expressed as the maximal-to-basal respiration ratio, and the respiratory control ratio was calculated as maximal respiration divided by proton leak. Parameters were derived from defined measurement phases: basal respiration was taken from the third measurement before injection, oligomycin-inhibited respiration as the minimum of measurements 4–6, FCCP-stimulated maximal respiration as the maximum of measurements 7–9, and non-mitochondrial respiration as the minimum of measurements 10–12.

To examine persistence of the respiratory changes, monocytes receiving a single mitochondrial dose of 5 μg of mitochondrial protein and untreated controls were assayed after 24 h or 6 days in culture, with five independent experiments per condition. These assays were not normalized to cell number. Accordingly, comparisons of respiratory coupling and relative reserve were restricted to ratios calculated within individual wells: maximal respiration/proton leak and maximal/basal respiration, respectively. For nuclear normalization in the separate cell-normalized experiment, nuclei were counted as follows. Immediately after the final measurement, the assay medium was removed, Hoechst 33342 was added to each well at 1:6,000 (v/v), and the plates were incubated for 10 min at 37 °C. We then imaged each well in the blue (DAPI) channel on an EVOS imaging system (Thermo Fisher Scientific) with a 2× objective, capturing the entire well in a single field; exposure and illumination settings were fixed and identical across all wells and plates. Nuclei were counted from these images in QuPath.

**Cell death analysis**

Cell viability and apoptosis were assessed using the Dead Cell Apoptosis Kit with Annexin V Alexa Fluor 488 and Propidium Iodide (Invitrogen, Thermo Fisher Scientific), according to the manufacturer's instructions.[2] Following 24 h of co-culture, untreated monocytes and MTM were harvested and washed with ice-cold Dulbecco's phosphate-buffered saline (DPBS) without calcium or magnesium. Cells were resuspended in 1× annexin V binding buffer containing 10 mM HEPES, 140 mM NaCl, and 2.5 mM $CaCl_2$ (pH 7.4). Alexa Fluor 488-conjugated annexin V and PI were added at the manufacturer's recommended volumes, using a 100 μg/mL PI working solution. Cells were incubated for 15 min at room temperature in the dark and subsequently analyzed using a five-laser Cytek Aurora spectral flow cytometer (Cytek Biosciences, Fremont, CA, USA). Viable cells were defined as annexin $V^{-}PI^{-}$, early apoptotic cells as annexin $V^{+}PI^{-}$, and membrane-compromised/dead cells as $PI^{+}$. Annexin $V^{+}PI^{+}$ cells encompass late apoptotic and necrotic populations, which cannot be distinguished conclusively using this assay alone.

**Cellular metabolic activity**

Cellular metabolic activity was assessed using Cell Counting Kit-8 (CCK-8; Dojindo Laboratories; Cat. No. CK04-01). Following 24 h of culture with or without transplanted mitochondria, 100 µL of undiluted CCK-8 reagent was added to 100 µL of culture medium per well, yielding a total assay volume of 200 µL. Samples were incubated for 2 h at 37 °C, and absorbance was measured at 450 nm using a SpectraMax® iD3 plate reader.

**THP1 Dual reporter assays**

THP1-Dual™ reporter cells (InvivoGen) were seeded in 96-well plates at 1 × $10^5$ cells per well. Cells carrying an NF-κB-inducible secreted embryonic alkaline phosphatase (SEAP) reporter and an interferon regulatory factor (IRF)-inducible Lucia luciferase reporter were exposed to 5 or 10 µg of isolated mitochondrial protein for 24 h. Unstimulated cells served as the reference condition, while Pam3CSK4 (10 ng/mL) and LPS ($10^3$ EU/mL) were included as stimulation controls. Culture supernatants were collected, and SEAP and Lucia luciferase activities were measured using QUANTI-Blue™ and QUANTI-Luc™ reagents (InvivoGen), respectively, according to the manufacturer's instructions. Absorbance and luminescence were recorded using a SpectraMax® iD3 plate reader. Reporter activities were expressed as fold induction relative to unstimulated cells. Four independent experiments were performed.

**Phagocytosis assay**

Cells and MTM was seeded in a 96 well plate at concentration of (1 x $10^5$ cells per well). To inhibit th eoxphos pathway, cells were pre-treated with rotenone (1 µM for 30 min at 37 °C) as negative control. Phagocytosis-associated fluorescence was assessed using pHrodo™ Deep Red *Escherichia coli* BioParticles™ (Thermo Fisher Scientific; Cat. No. P35360). Following 24 h of culture with or without transplanted mitochondria, the culture medium was removed, and 100 µL of a particle suspension at 1 mg/mL resuspended in PBS was added to each well, providing 100 µg of particles per well. Monocytes and MTM were incubated with the particles for 2 h at 37 °C. Following incubation, cells were analyzed using an Attune CytPix flow cytometer (Thermo Fisher Scientific). The pHrodo Deep Red signal was quantified as median fluorescence intensity (MdFI) within the gated monocyte population.

**Statistical analysis**

Data were analyzed using GraphPad Prism 11. The biological replicate was an individual healthy donor for primary monocyte experiments, an independent mitochondrial preparation for isolated mitochondrial assays, and an independent experiment for THP1-Dual reporter assays. Technical replicates were averaged within each biological replicate and treatment condition before statistical analysis. For cell-normalized Seahorse experiments, normalized measurements from the two plates were combined within donor and treatment condition, yielding one value per donor per condition. Donor matching was retained in comparisons between treatment conditions. Matched comparisons were evaluated using paired tests or repeated-measures analyses, as specified for each figure. Isolated mitochondrial OCR was analyzed after logarithmic transformation using repeated-measures one-way ANOVA with the Geisser–Greenhouse correction. Surface-marker fold changes were calculated from median fluorescence intensity (MdFI) and compared by ratio paired *t*-tests on log-transformed values, with geometric means and 95% confidence intervals back-transformed for presentation. Benjamini–Hochberg adjustment was applied across the six surface markers within each comparison and is reported alongside the effect sizes in Supplementary Figure 6. Donor-matched respiratory and cytokine fold changes were evaluated on the logarithmic scale, and estimates and 95% confidence intervals were back-transformed for presentation.

ANOVA comparisons used Tukey's, Dunnett's or Šídák's multiple-comparison tests, as specified in the corresponding figure legends. Donor-matched fold changes were compared by ratio paired t-tests on log-transformed values. Where a marker's compensated condition mean fell at or below zero in a donor, that donor was excluded from the fold-change comparison for that marker, giving n = 5 for CD16 and n = 4 for CD163 in the MITO comparison and n = 2 for CD163 in the antimycin A comparisons. Tests were two-tailed except for TNF-α and IL-10 in the LPS response, for which a one-tailed test was used on the basis of a direction specified before analysis; the lack of significance under two-tailed testing is also noted for IL-10. No correction was made for multiplicity across the 13 analytes. Data presentation, biological replicate numbers and the statistical test used are specified for each experiment. A P value below 0.05 was considered statistically significant.

**Supplementary Figures**

**Supplementary Figure 1**

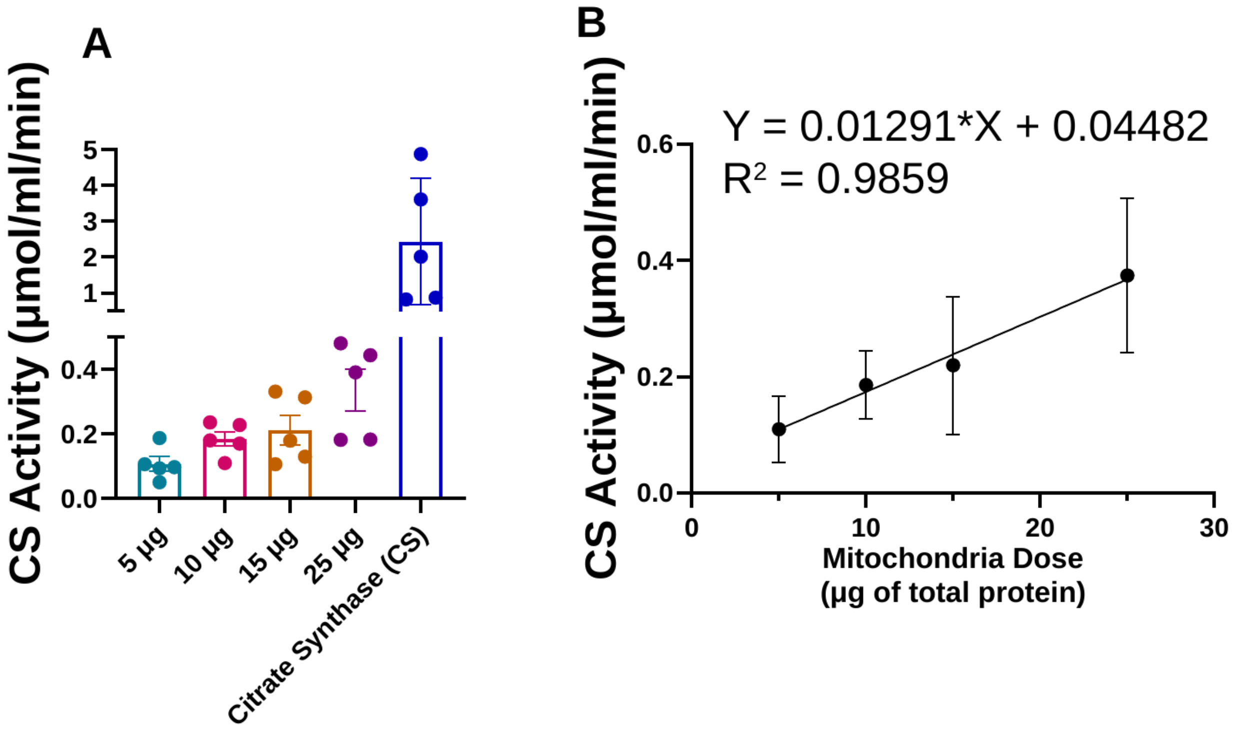


**Supplementary Figure 1. Citrate synthase activity scales with mitochondrial protein content.** (A) Citrate synthase activity of isolated mitochondria at 5, 10, 15 and 25 µg of mitochondrial protein, with protein content determined by Bradford assay, with purified citrate synthase (CS) assayed alongside as a positive control. Bars are mean ± SEM of 4 to 5 replicates per dose. (B) Linear regression of mean citrate synthase activity on mitochondrial protein, with SD at each dose: Y = 0.0129X + 0.0448, $R^2$ = 0.986, slope 95% CI 0.0083–0.0176, P = 0.007 across four doses. The intercept (0.045, 95% CI −0.029 to 0.118) does not differ from zero, consistent with activity being proportional to protein. Citrate synthase, a matrix marker enzyme, was therefore used alongside protein content to confirm that protein mass tracks mitochondrial content in these preparations; doses are given as µg of mitochondrial protein throughout. Activity is expressed in µmol/ml/min.

### Supplementary Figure 2

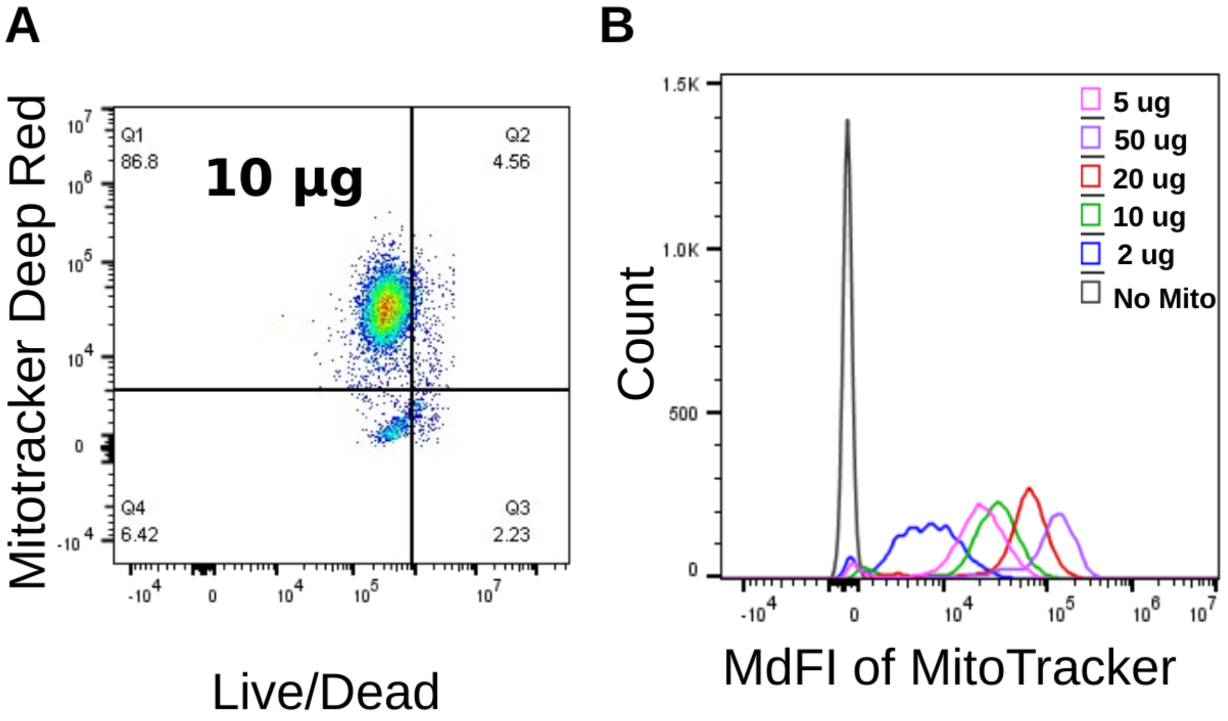


**Supplementary Figure 2. Dose-dependent uptake of isolated mitochondria by human monocytes with preserved viability.** Monocytes (2.3 × $10^5$ per well) were cultured for 24 h with isolated mitochondria pre-labeled with MitoTracker Deep Red at 2, 5, 10, 20 or 50 µg of mitochondrial protein per well, or without mitochondria (No Mito), and then analyzed by flow cytometry. (A) Representative dot plot of MitoTracker Deep Red fluorescence (y axis) against viability dye (x axis) at 10 µg; quadrant gates were set on the no-mitochondria control. (B) Overlaid histograms of MitoTracker Deep Red fluorescence in viable monocytes at each dose, with unlabeled cells (No Mito) as the reference. Fluorescence increased with dose while the proportion of viable cells was unchanged across the range tested. The 5 µg dose used throughout the study is also shown in Figure 2B.

**Supplementary Figure 3**

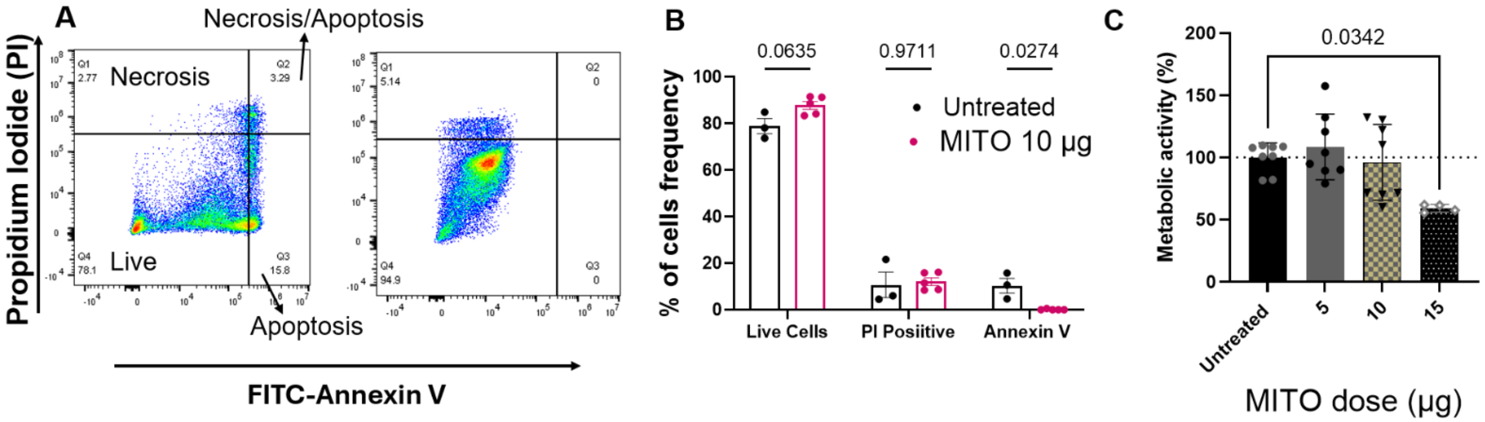


**Supplementary Figure 3. Monocyte viability and metabolic activity after mitochondrial transplantation.** Monocytes (2.3 × $10^5$ per well) were cultured for 24 h alone or with isolated mitochondria, then stained with Alexa Fluor 488-conjugated annexin V and propidium iodide (PI) and analyzed by flow cytometry. (A) Representative dot plots of annexin V–Alexa Fluor 488 (x axis) against PI (y axis) for untreated monocytes and after transplantation of 10 μg. Quadrants: live (annexin V−PI−, lower left), apoptotic (annexin V+PI−, lower right), membrane-compromised (annexin V−PI+, upper left) and late apoptotic or necrotic (annexin V+PI+, upper right). (B) Percentage of cells in each quadrant. The annexin V-positive fraction was lower after transplantation than in untreated monocytes (P = 0.027); the live and PI-positive fractions did not change. (C) Metabolic activity measured by CCK-8 as absorbance at 450 nm, expressed as a percentage of untreated monocytes. Activity was maintained at 5 and 10 μg and fell at 15 μg (P = 0.034). Bars are mean ± SEM. (B) Two-way ANOVA followed by Šídák's multiple comparisons test, n = 3 to 5. (C) One-way ANOVA with Tukey's multiple comparisons test, n = 4.

**Supplementary Figure 4**

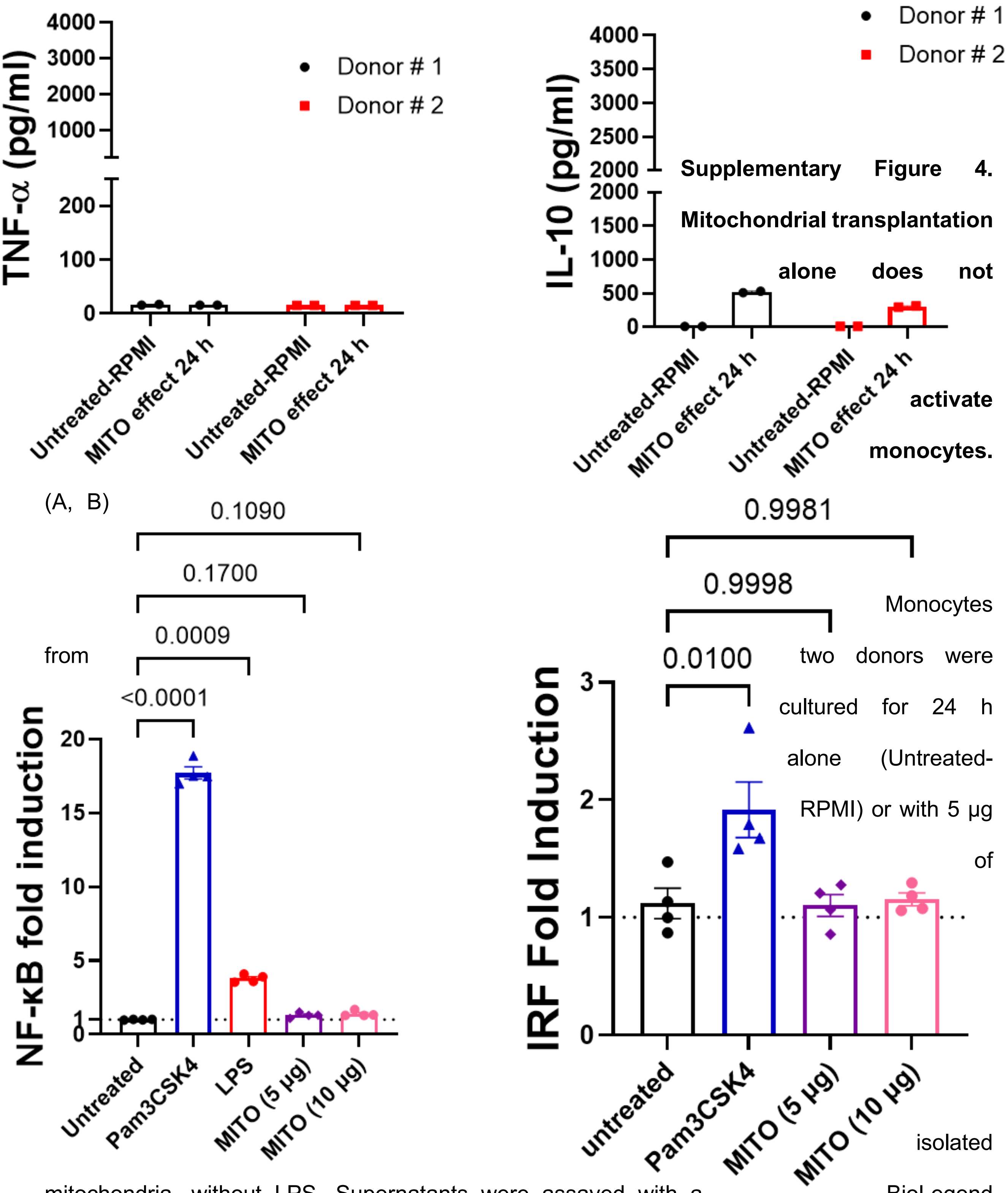


**Supplementary Figure 4. Mitochondrial transplantation alone does not activate monocytes.** (A, B) Monocytes from two donors were cultured for 24 h alone (Untreated-RPMI) or with 5 µg of isolated mitochondria, without LPS. Supernatants were assayed with a BioLegend

LEGENDplex bead-based immunoassay. Neither TNF-α (A) nor IL-10 (B) rose above the untreated level in either donor. (C, D) THP1-Dual reporter cells, which carry an NF-κB-inducible SEAP reporter and an IRF-inducible Lucia luciferase reporter, were exposed for 24 h to 5 or 10 μg of isolated mitochondria. NF-κB activity (C), read as QUANTI-Blue absorbance, and IRF activity (D), read as QUANTI-Luc luminescence, are expressed as fold induction over unstimulated cells. Pam3CSK4 (10 ng/ml) raised NF-κB 17.6-fold ($P < 0.0001$) and LPS ($10^3$ EU/ml) raised it 4-fold ($P = 0.0009$); Pam3CSK4 raised IRF 1.9-fold ($P = 0.010$). Mitochondria induced neither reporter at either dose (NF-κB, $P = 0.17$ and 0.11; IRF, $P > 0.99$ for both). Each point is one donor in (A, B; $n = 2$) and one independent experiment in (C, D; $n = 4$); bars are mean ± SEM. (C, D) One-way ANOVA with Tukey's multiple comparisons test.

**Supplementary Figure 5**

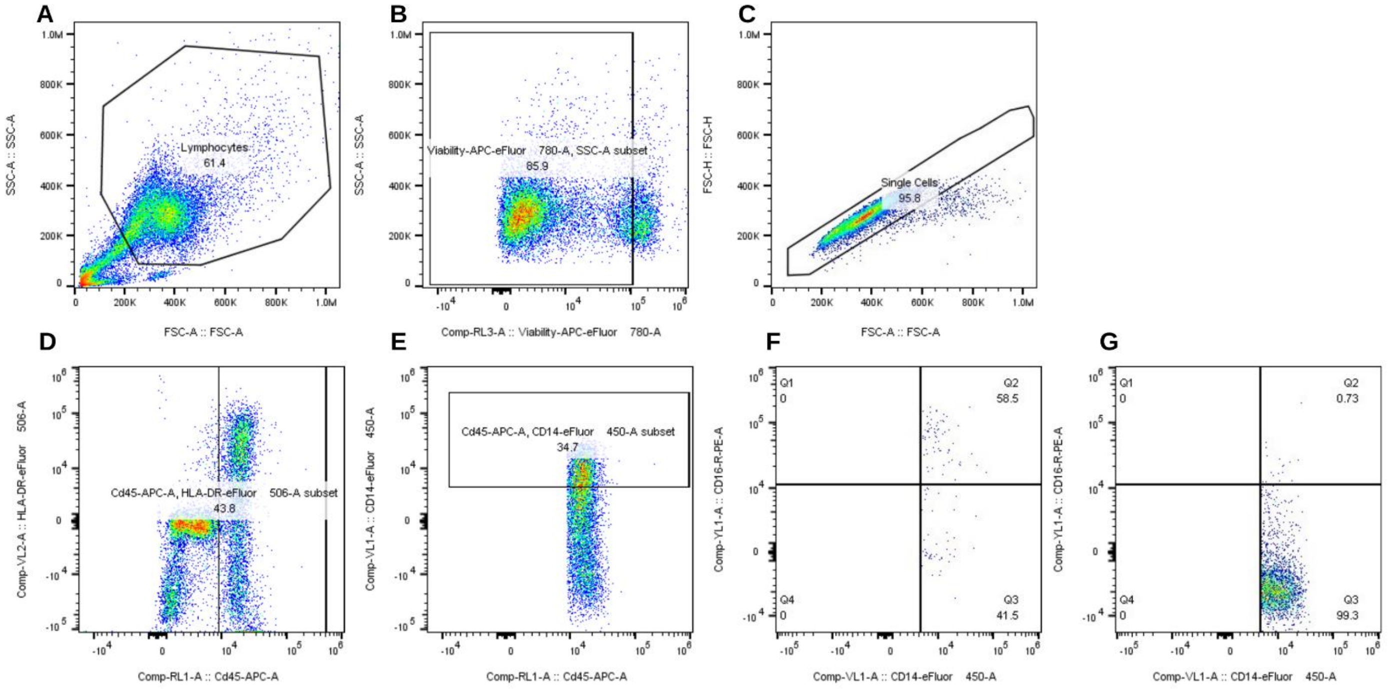


**Supplementary Figure 5. Gating strategy for monocyte subsets and surface markers.** Representative gating from one donor. (A) Cells were identified on SSC-A against FSC-A, excluding debris. (B) Dead cells were excluded with fixable viability dye. (C) Single cells were selected on FSC-H against FSC-A. (D) HLA-DR against CD45 identified HLA-DR-positive leukocytes, reducing contributions from HLA-DR-negative T cells, NK cells, and granulocytes. (E) CD14 against CD45 gated CD14-positive monocytes. (F, G) CD14 against CD16 resolved the classical (CD14++CD16−), intermediate (CD14++CD16+) and non-classical (CD14+CD16++) subsets, with quadrants set on fluorescence-minus-one (FMO) controls for CD14 and CD16.

### Supplementary Figure 6

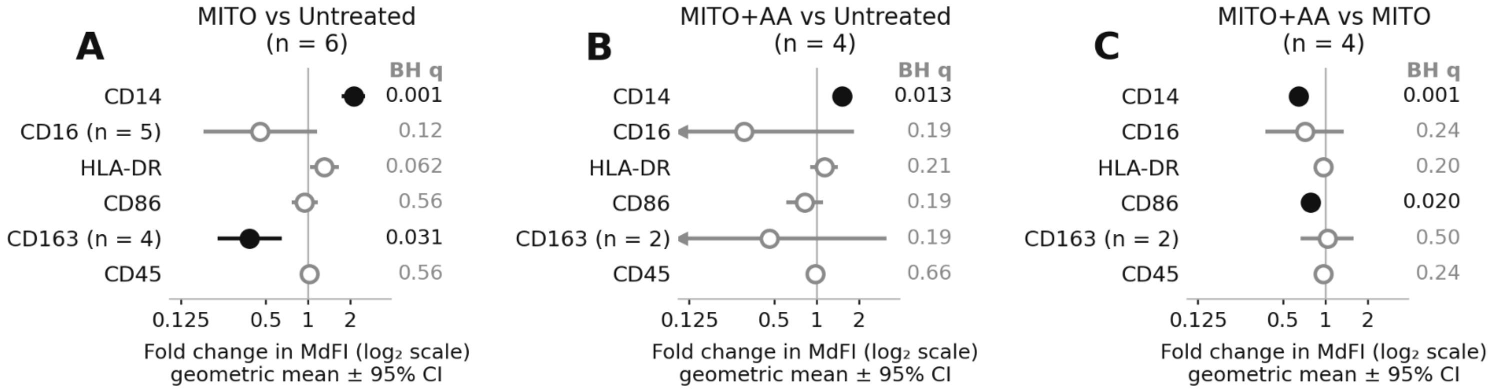


**Supplementary Figure 6. Effect sizes and multiplicity-adjusted significance for all six surface markers.** Monocytes were cultured for 24 h alone (Untreated), with 5 µg of isolated mitochondria (MITO), or with antimycin A-pretreated mitochondria (MITO+AA). Each panel shows one comparison: (A) MITO versus Untreated (n = 6 donors), (B) MITO+AA versus Untreated (n = 4) and (C) MITO+AA versus MITO (n = 4). Points are the geometric mean fold change with 95% confidence intervals on a $\log_2$ axis; the vertical line at 1 marks no change, arrowheads mark intervals that extend beyond the axis, and filled symbols mark the markers that remain significant after Benjamini–Hochberg adjustment across the six markers within each comparison (q values shown at right). Estimates come from the same ratio paired t-tests as Figure 3C–E and use the same donors: where compensation left a marker's condition mean at or below zero in a donor, that donor is excluded from that marker and the reduced n is given beside the marker name. The direction in (C) is opposite to that described for Figure 3E: a value below 1 here indicates lower expression with antimycin A-treated mitochondria than with respiration-competent mitochondria.

### Supplementary Figure 7

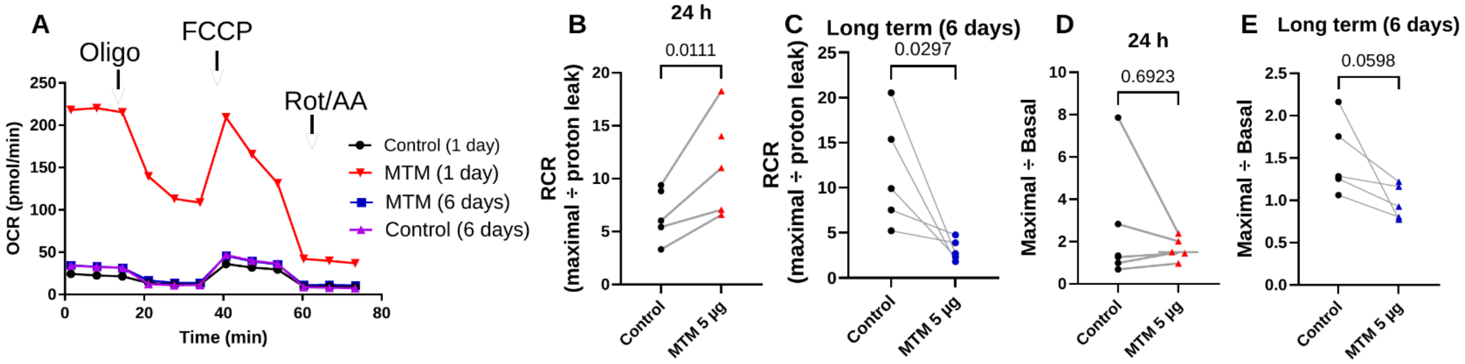


**Supplementary Figure 7. Respiratory coupling after mitochondrial transplantation at 24 h and after 6 days in culture.** Monocytes received a single 5 µg dose of isolated mitochondria (MTM) or were left untreated (Control), and were assayed with the Seahorse XF Cell Mito Stress Test after 24 h or after 6 days in culture, to ask whether the change persists in extended culture. Rates are per well and are not normalized to cell number, so panels B–E show ratios taken within each well, which reduce sensitivity to cell number but do not control for changes in viability or cell composition. (A) OCR over time for Control and MTM at each culture duration; arrows mark the injection of oligomycin, FCCP and rotenone/antimycin A. Mean ± SEM. (B, C) Respiratory control ratio (maximal ÷ proton leak) at 24 h and at 6 days. RCR rose at 24 h (1.74-fold, 95% CI 1.27–2.39, P = 0.011; higher in 4 of 4 experiments) and fell by 6 days (0.28-fold, 95% CI 0.10–0.81, P = 0.030; lower in 5 of 5), where proton leak was 15.7-fold higher than control (95% CI 4.1–60.8, P = 0.005). (D, E) Maximal ÷ basal respiration at 24 h (0.88-fold, 95% CI 0.38–2.02, P = 0.69) and at 6 days (0.66-fold, 95% CI 0.42–1.03, P = 0.060; lower in 5 of 5). FCCP failed to raise OCR above baseline in 4 of 5 MTM wells at 6 days. Each point is one independent experiment; lines join Control and MTM on the same plate; bars are means. n = 5 experiments per arm, except RCR at 24 h (n = 4; one control plate had a proton leak at or below zero, leaving RCR undefined). P values are from ratio paired t-tests on the experiment values and are unadjusted.

**Supplementary Figure 8**

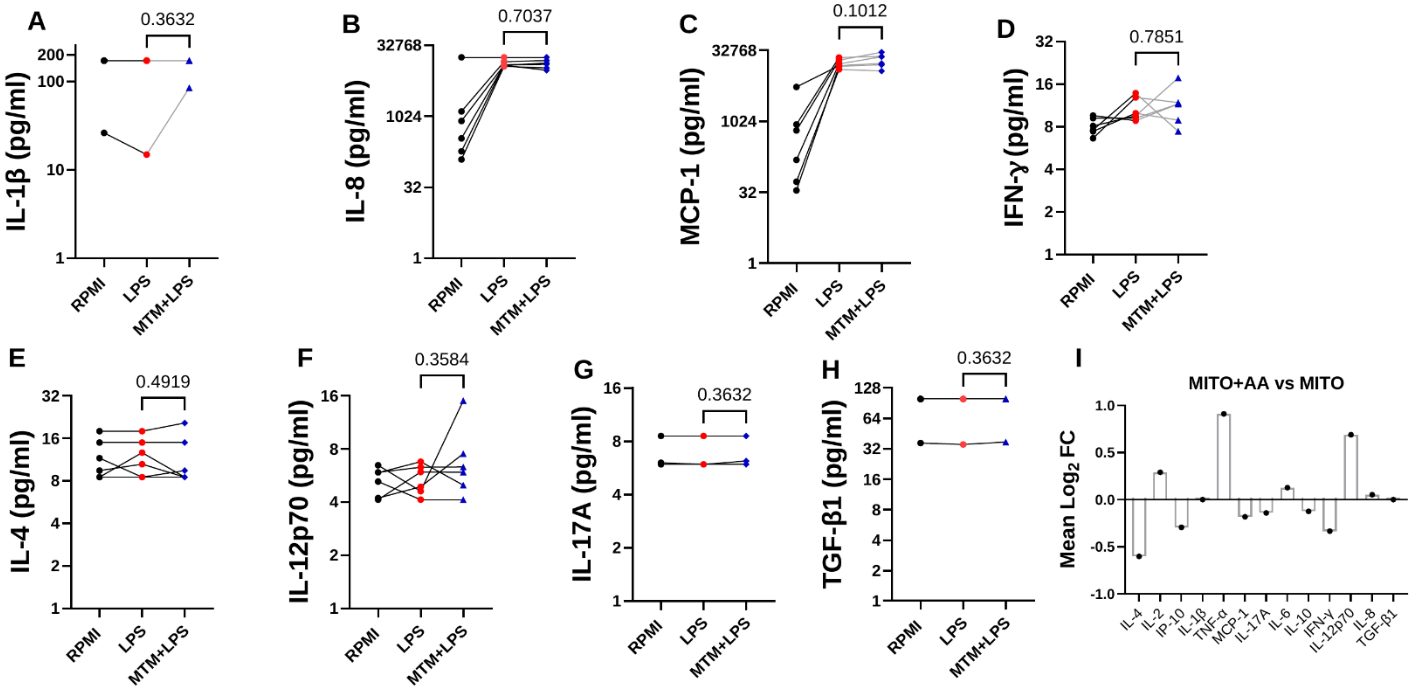


**Supplementary Figure 8. Remaining cytokines and chemokines in the LPS response of transplanted monocytes.** Monocytes were co-cultured for 24 h with 5 µg of isolated HEK293T mitochondria, then stimulated with LPS ($10^3$ EU/ml) for 24 h. Supernatant analytes were quantified by bead-based multiplex immunoassay (LEGENDplex). (A) IL-1β (P = 0.3632), (B) IL-8/CXCL8 (P = 0.7037), (C) MCP-1/CCL2 (P = 0.1012), (D) IFN-γ (P = 0.7851), (E) IL-4 (P = 0.4919), (F) IL-12p70 (P = 0.3584), (G) IL-17A (P = 0.3632) and (H) TGF-β1 (P = 0.3632) in RPMI, after LPS alone and after LPS in transplanted monocytes (MTM+LPS); each point is one donor and lines join the same donor. n = 6 donors, ratio paired t-test on donor-matched fold changes, two-tailed. IL-1β, IL-17A and TGF-β1 were below the assay limit of quantification in 5 of 6 donors, so these comparisons are uninformative and the P values reflect that structure rather than an absence of biological effect. (I) Mean $\log_2$ fold change in monocytes given antimycin A-treated versus respiration-competent mitochondria, across all analytes, to explore differences between the two preparations; n = 3 donors, bars are mean ± SEM. Antimycin A pretreatment was associated with a higher TNF-α point estimate (+0.91 $\log_2$) and a slightly lower IL-10 point estimate (−0.12 $\log_2$), although no individual analyte reached significance at this sample size ($P \geq 0.18$).

**Supplementary Figure 9**

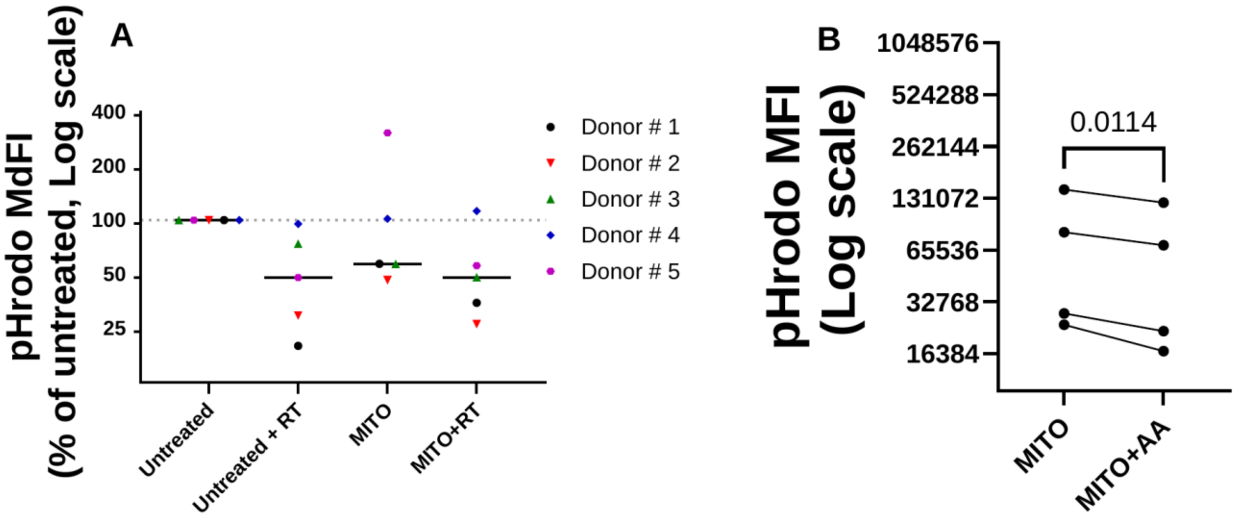


**Supplementary Figure 9. Phagocytosis-associated fluorescence after mitochondrial transplantation.** Monocytes were cultured for 24 h alone (Untreated) or with 5 µg of isolated mitochondria (MITO), then incubated with pHrodo-labeled particles, which fluoresce as the phagosome acidifies. Where indicated, cells were pretreated with 1 µM rotenone (RT) for 30 min before the assay. (A) pHrodo MdFI as a percentage of each donor's own untreated cells in five donors; bars are medians. Rotenone lowered uptake in untreated monocytes to a median of 48% (range 20–95%; geometric mean 0.46-fold, P = 0.052). Transplantation alone gave a median of 57% (range 46–306%), but the change across donors was not significant (0.86-fold, P = 0.68): uptake fell in four donors and rose about threefold in donor 5. Uptake after transplantation remained rotenone-sensitive (MITO+RT, median 48%; 0.49-fold versus untreated, P = 0.043). (B) pHrodo MdFI in monocytes given respiration-competent (MITO) or antimycin A-pretreated (MITO+AA) mitochondria, in the four donors assayed with both; lines join the same donor. Uptake was 1.27-fold higher with respiration-competent mitochondria (95% CI 1.11–1.45, P = 0.011; higher in 4 of 4 donors). Ratio paired t-tests on donor ratios; each point is one donor, and the same donor numbering applies to both panels.
